# Adolescent pragmatic development mirrors pragmatic individual differences in adulthood: An fMRI study

**DOI:** 10.1101/2025.06.27.661903

**Authors:** Christoffer Forbes Schieche, Julia Uddén

## Abstract

Pragmatic development is still ongoing during adolescence. However, the developmental trajectory during this period is understudied and the underlying neural correlates of pragmatic development are unknown. In this study, we used an established fMRI paradigm contrasting indirect and direct speech acts in a newly acquired adolescent sample (n = 51) and a previously collected adult sample (n = 57). The adolescent sample was subdivided into Young (ages 13-15) and Mid (ages 16-18) groups and the adult sample (Old, ages 20-36) was split based on pragmatic skill level (established in separate behavioral tests). We observed age-related increases in activity in the posterior medial superior frontal gyrus, and a combination of age-related differences and adult individual differences in the dorsal aspects of the posterior cingulate cortex (dPCC). Both clusters were located outside the classic perisylvian language network. We interpret the dPCC findings in terms of its overlap with the Default Mode Network. Similar activity in the dPCC between adolescents and adults with lower pragmatic skill is consistent with a view that a delayed adolescent development may lead to persistent pragmatic difficulties in adulthood. The findings also indicate development towards context-specific processing. By studying adolescence, a presumably uniquely human extended developmental period, we provide a rare empirical angle on the question of which aspects of brain and cultural evolution contributed to the human communicative faculty.

## 1. Introduction

Successful everyday communication requires more than retrieval of words or the application of syntactic rules. It also involves continuous interpretation beyond the literal content of a message, to understand others’ communicative intentions. Such pragmatic processes encompass a range of phenomena in comprehension, such as speech act interpretation and the integration of multimodal signals. Several aspects of communicative production and interaction—such as audience design and turn-taking—are also pragmatic in nature. However, in this article we will primarily focus on pragmatic comprehension.

Recent evolutionary accounts suggest that pragmatics plays a key role in what makes us human, enabling complex and flexible interaction on a level not observed in other species (Levinson, 2006). Based on studying communicative situations with infants and chimpanzees, it has been suggested that the uniquely human language faculty is rooted in evolved social and communicative contexts, which have provided a foundation for language use to emerge (M. Tomasello et al., 2012). Moreover, while claims of linguistic universals at the level of lexical and grammatical meaning have been contested (Evans & Levinson, 2009), cross-linguistic evidence of recurring pragmatic and conversational patterns (Dingemanse et al., 2013, 2015) highlights a potentially important role for pragmatics in human communication and its emergence.

Supporting evidence for the unique role of pragmatics in the human lineage emerges already in early development. Infants are sensitive to ostensive communicative behaviors and signals, including eye-contact and infant-directed speech (Cooper & Aslin, 1990; Farroni et al., 2002), from the first months of life. By six months, they tend to track others’ gaze when these signals occur (Senju & Csibra, 2008). Infants show increased attention and inferential behaviors when exposed to patterns that more closely resemble human communication (Movellan & Watson, 2002; Tauzin & Gergely, 2018) and young children (2-2.5 years) outperform both age-matched and adult non-human primates in areas involving social understanding and cooperation specifically (Herrmann et al., 2007; Wobber et al., 2014).

Another trait often considered uniquely human is an extended childhood. Compared to other species, including our closest primate relatives, the period from human infancy to independence (i.e., adulthood) is significantly longer (Bogin, 1990, 1997). Among the proposed evolutionary advantages of this extended childhood are that it allows for continuous learning of cultural and other complex knowledge (Bogin, 1990, 1997; Uomini et al., 2020). Notably, the adolescent phase may be especially crucial for social learning and exploration (Gopnik, 2020; Gopnik et al., 2017). Abilities related to pragmatics, such as social cognition and Theory of Mind (ToM), also appear to undergo continued development during this period (Bosco et al., 2014; Brizio et al., 2015). This is further supported by observations of developmental changes in brain activity related to social processing and reasoning during adolescence (Crone & Dahl, 2012; Dumontheil, 2015).

This paper builds on the idea to combine these observations of uniquely human (1) pragmatic abilities and (2) extended childhood. What if pragmatics is one of the core abilities learned or refined during this extended childhood—contributing to what makes us uniquely human? This question invites investigation into the function of brain structures and brain networks showing accelerated evolution in humans and continued development during adolescence (Arain et al., 2013; Blakemore, 2012; Blakemore et al., 2007; Catani & Bambini, 2014; Crone & Dahl, 2012; Dahl, 2004; Dumontheil, 2015; Giedd et al., 1999; Stiles & Jernigan, 2010). To the extent that neural development supports pragmatic functions and indexes their maturation, we can not only begin to understand *what* makes us uniquely human, but also how this came about, by focusing on relevant aspects of brain evolution and development.

Considering the existing literature on pragmatic development, there is a relative gap in studies investigating the adolescent period, as most developmental accounts have focused on early childhood and the pre-adolescent period (e.g., Matthews et al., 2018). Nonetheless, both observational and experimental work support a general assumption of a continuous development of pragmatics during adolescence (Arvidsson et al., 2022; Garraffa & Mazzaggio, 2025; Nippold, 1993, 2000; Petit et al., 2025). Raffaelli and Duckett (1989) observed an increase in the sheer duration spent in conversation with peers in mid-adolescence, and the amount of distinct conversational exchanges—especially non-familial ones—also increases (Nippold, 1993). Successful navigation of such diverse conversational contexts demands increasingly flexible and sophisticated conversational skills. More recently, experimental work has provided evidence of increasing pragmatic maturity from early into mid-adolescence in pragmatic production (Arvidsson et al., 2022). Petit et al. (2025) have also demonstrated a continuous increase in six different pragmatic comprehension and production measures, from adolescence into early adulthood (peaking ages 25-30). It should be noted that none of the mentioned studies have examined the pragmatic phenomenon operationalized in the present study, i.e. indirect speech acts (see following paragraphs). Moreover, there are currently no published neuroimaging studies directly addressing pragmatic development in adolescence (see however preprint by Asaridou et al., 2019).

To investigate the development of pragmatics in adolescence, we here operationalize pragmatic processing through indirect speech acts (ISA). In ISA, unlike direct speech acts, the literal utterance and intended meaning do not match and the properties of ISA and how they are understood have long been discussed within theoretical pragmatics (Grice, 1975; Searle, 1975). In line with such perspectives, understanding the intended meaning of an ISA requires making inferences based on contextual information, which can include common world knowledge or considering the relation to the specifics of the conversation. Consider the utterance “She is not so used to Asian seasoning” as a reply to (1) “Why doesn’t your girlfriend like Japanese food?” or (2) “Did your girlfriend like my vegan noodles?”. As a response to (1), the reply provides the requested information, whereas in (2) the requested information (yes/no) is not directly given. However, (2) would typically be understood as indicating that the girlfriend disliked the noodles. Furthermore, using indirectness is often linked to politeness and management of one’s own and others’ face (Brown & Levinson, 1987; Goffman, 1955). A direct expression of the intended negative meaning in (2) could be considered impolite and constitute a face-threatening act: paraphrasing the criticism in an indirect manner functions instead as a face-saving act. The differences between (1) and (2), in terms of inferential demands beyond the literal meaning and face considerations, suggest that the comprehension of these two types of utterances may involve different processing demands. As will be described in later sections, in the present study we used direct informative replies and indirect face-saving criticisms like the ones above.

Within the adult population, multiple fMRI studies have examined the cognitive architecture underlying pragmatic abilities by contrasting different types of speech acts (e.g., statements, questions, requests) (Bašnáková, 2019; Bašnáková et al., 2014; Bendtz et al., 2022; Egorova et al., 2016; Feng et al., 2021; Hellbernd & Sammler, 2016, 2018; van Ackeren et al., 2016; see also review by R. Tomasello et al., 2025). A number of these have specifically contrasted indirect speech acts (ISA) with direct speech acts (Bašnáková et al., 2014; Bendtz et al., 2022; Feng et al., 2021; van Ackeren et al., 2016). These studies have consistently shown increased activation within the canonical language (Fedorenko et al., 2024; Hagoort, 2017; Hickok, 2009) and Theory of Mind (ToM) (Schurz et al., 2014) networks during the processing of ISA. This suggests that the processing of ISA engages both structural language processes and ToM-related abilities (such as perspective taking) to a greater extent than direct speech acts. However, findings from our previous study (Bendtz et al., 2022) indicate that these two faculties do not fully account for pragmatic processing. In that study, adult participants were divided into two groups based on behavioral tests of pragmatic production and comprehension. The groups consisted of those with the lowest scores on both measures (the LS group) and those with the highest scores (the HS group). When we compared the two groups in a test for interaction (HS > LS, Indirect > Direct), we observed higher activity for the HS group in two clusters outside of the language and ToM networks: (1) the left anterior/mid intraparietal sulcus and (2) the bilateral dorsal precuneus (together referred to as the ParPrec clusters). This led us to conclude that pragmatics is at least partially segregable from structural language and ToM processes, a conclusion that we have further strengthened in a recent investigation using functional connectivity measures (Forbes Schieche et al., 2025).

### The current study

While previous fMRI research provide insights into the neural correlates of pragmatic processing, this body of literature is so far limited to adult participants. The developmental period of adolescence is yet to be investigated from a neural perspective. In this study, we thus investigated the neural basis and developmental trajectory of indirect speech act (ISA) processing during adolescence into early adulthood. We used a newly acquired sample of adolescents, further subdivided into two groups based on age (Young: ages 13—15 and Mid: ages 16—18), as well as our previously collected sample from Bendtz et al. (2022) (here named the Old group: ages 20—36). By using the same ISA paradigm in both samples (Bendtz et al., 2022) we could compare the groups directly. Although adolescent pragmatic development remains underexplored, the paradigm presents several different conversations (see 2.3), parallelling the increase in conversational experiences during adolescence.

Our investigation was guided by four research questions (defined in our preregistration and outlined below). To address these questions, we first examined whole-brain activation during pragmatic processing in adolescents (RQ1). Next, we investigated age-related differences using both whole-brain and ROI approaches (RQ2–3). Finally, we combined whole-brain and ROI analyses to explore whether delayed development could explain individual differences in adult pragmatic skill (RQ4). In our ROI analyses, we used both predefined ROIs and ROIs derived from the data. While Bendtz et al. (2022) focused on the Indirect > Direct contrast, we also examined the reverse (Direct > Indirect) for completeness.

#### RQ1. What is the neural basis of pragmatic processing in adolescents?

First, we examined whole-brain activation for indirect versus direct speech acts (Indirect > Direct) within adolescents. We hypothesized that activation during pragmatic processing (ISA) would resemble earlier findings from adults (Bašnáková et al., 2014; Bendtz et al., 2022), i.e., that the language network and additional regions (such as the medial superior frontal gyrus (mSFG)/medial prefrontal cortex (mPFC)) would be more active during ISA.

#### RQ2-3. What is the developmental trajectory of pragmatic processing within adolescence and from adolescence to adulthood?

While we expected the general pattern observed in adults to partially replicate in adolescents, our main interest was to test potential age-related differences statistically. These tests were performed between the adolescent groups (RQ2) and between the adolescents and adults (RQ3). Based on prior pragmatic (Asaridou et al., 2019; Bendtz et al., 2022) and related developmental literature (social cognition/ToM, Blakemore et al., 2007; social emotion, Burnett et al., 2009), we hypothesized an anterior-to-posterior functional shift across the three age groups (Young, Mid, and Old), which we directly tested using three predefined ROIs. We expected lower activity within the anterior mSFG (amSFG) in the adults relative to the adolescents. More posteriorly, we hypothesized that activity within the posterior mSFG (pmSFG) would instead increase across the three age groups, based on the lack of activity in this region for adolescents (Asaridou et al., 2019) but its presence in adults (Bendtz et al., 2022). Similarly, we expected an increase in activity across age groups in the anterior temporal lobe (ATL), which is posterior relative to the amSFG. Altogether, such a pattern is thus what we consider a developmental posterior shift.

#### RQ4. Can challenges in pragmatic skill in adulthood be explained by delayed development?

For our final research question, we explored whether adult pragmatic challenges may partly reflect delayed development. Here we made use of the two groups from Bendtz et al. (2022) (i.e., the HS and LS groups). The participants of the LS group were selected from those with the lowest scores on two behavioral pragmatic tests; as such, they could be considered to have challenges in their pragmatic skill compared to a normative adult. We hypothesized that they would show brain activity that reflects less developed or mature processing. We therefore statistically examined similarities and differences between adolescents and the adult groups to test whether individual variation in pragmatic skill may relate to developmental trajectories (see pre-specified testing procedure in section 2.6.3 in Methods).

In summary, the current study expands the scope of neuropragmatic research by examining the neural basis and developmental trajectory of pragmatic processing during adolescence. This developmental stage is uniquely extended in humans, allowing for the continued refinement of complex communicative abilities, such as pragmatics. By contrasting younger and older adolescents with adults we capture how such communicative capacities mature neurally. Finally, by comparing these trajectories to adult individual variation in pragmatic ability, we provide a foundation for understanding both typical development and what mechanisms may underlie pragmatic challenges in adulthood.

## 2. Method

### 2.1 Participants

55 participants (excluding six pilot participants) were invited to take part in the experiment in the MR-scanner (32 females, ages 13—18, mean age 15.5 ± 1.7). These were further subdivided into two age groups: the Young group aged 13—15 (n = 27, 13 females, mean age 13.9 ± 0.8) and the Mid group aged 16—18 (n = 28, 19 females, mean age 17.0 ± 0.7). Criteria for participation included that participants had to be native Swedish speakers (or native level at seven years old), no history of substance abuse, neurological impairment, brain surgery, ADHD or ASD diagnosis, or language impairment. Each participant received gift cards valued SEK 297 after completing the experiment. All data was acquired at Stockholm University Brain Imaging Center (SUBIC) and the data collection was approved by Swedish Ethical Review Authority. All participants gave informed consent, as well as their guardians in case they were younger than 15. Due to exclusion criteria, four participants were excluded from the final analysis, resulting in a final sample of n = 51 (Young, n = 24 (22 right-handed, self-reported), and Mid, n = 27 (25 right-handed)). Some individual runs were also excluded. For details on the basis for exclusion and exclusion criteria from data quality, see section 2.4.2 and Supplementary Materials §S2.1.

The adult sample (again, referred to as the Old group) consisted of 57 participants (age 20—36, 31 females, no handedness data available), and was collected and analyzed in a previous investigation within our lab (Bendtz et al., 2022). The sample can be further subdivided into two groups based on pragmatic ability measured through two behavioral tests at a prior date to acquisition of the MR data. Participants with the worst performance on both tests form a low-scoring group, LS (n = 28, 13 females, mean age 29.6 ± 4.9), and those with the highest performance form a high-scoring group, HS (n = 29, 18 females, mean age 30.1 ± 4.1).

### 2.2 Preregistration

Our research questions, hypotheses, and analysis plan were preregistered on Open Science Framework (OSF) prior to performing the final analysis (https://osf.io/ft9cp/files/kb96c). Note that the ROIs we here call the anterior and posterior medial superior frontal gyri (amSFG/pmSFG, see section 2.6) were named the anterior and posterior medial prefrontal cortex (amPFC/pmPFC) in the preregistration.

### 2.3 fMRI paradigm and stimulus

We used the same paradigm as in Bendtz et al. (2022), with slight modifications. The paradigm consists of an event-related auditory design with several conversations between interlocutors, presented in Swedish. All conversations include an introductory context (explaining a situation), a question from one of the interlocutors, which the other one answers either in a direct or indirect manner, i.e., performs a direct or indirect speech act. The main interest of the paradigm is the answers which exist in pairs that are linguistically identical but rendered direct or indirect based on varying the question posed by one of the interlocutors. In some cases, the context was slightly altered to better facilitate either condition. No changes were made to the conversations, as the content was considered suitable also for adolescent participants. Each participant heard only one version of each trial.

All the direct speech acts were informative replies whereas all indirect speech acts served a face-saving function (Brown & Levinson, 1987). Behavioral studies have shown that the interpretation of indirect speech acts as such is more obvious when they are posed in a face-saving context (Holtgraves, 1998). Furthermore, the indirect speech acts conveyed a critical opinion or negative message directed at the interlocutor: in half of the trials, the opinion belonged to the speaker, and in the other half it belonged to a third-party not present in the conversation.

In total, there were 78 experimental trials, each in a direct and indirect version. Each participant heard 39 indirect and 39 direct trials throughout the experiment. The experiment also included 14 filler trials and 12 control questions. The filler trials had the same structure as the experimental trials but ended with a direct yes/no reply to a question. The 12 control questions were paired with particular trials (regardless of direct or indirect). After these trials, a yes/no question was (auditorily) posed to the participant about some detail regarding the conversation they had just listened to. Participants replied to the question with a button box. Some of the control questions were changed from Bendtz et al. (2022), see Supplementary Materials §S2.3 for more details. Table 1 shows examples of each type of trial. The experiment was divided into three runs, each lasting between 10—12 minutes. Each run contained 26 experimental trials (13 direct, 13 indirect), 4—5 filler trials, and 3—5 compliance questions. Further information about trial order can be found in Supplementary Materials §S2.3.1.

**Table 1.** Examples of each trial type used in the paradigm, translated from Swedish.

| Table 2: Examples of each trial type used in the paradigm, translated from Swedish. |  |  |  |
| --- | --- | --- | --- |
| Trial type | Context | Question | Answer |
| Direct | Magnus and Emilia are old friends. They are discussing how hard it is to find restaurants which both you and your partner fancy. Emilia asks Magnus: | Why doesn't your girlfriend like Japanese food? | She is not so used to Asian seasoning. |
| Indirect | Magnus and Emilia are old friends. They are talking about the last time Magnus visited Emilia in her student dormitory. Emilia asks Magnus: | Did your girlfriend like my vegan noodles? |  |
| Control question | Was there someone who was not so used to Asian seasoning? |  |  |
| Filler | Benny and Ellinor are doing their laundry. Benny went out walking the dog but is now back. He asks Ellinor, who was the last to visit the laundry room: | Is the washing machine done? | Yes, just put everything in the dryer. |

### 2.4 fMRI data acquisition

This section describes acquisition and preprocessing specific to the adolescent data. The pipeline was close to identical to the one used for the adult data set, except for the settings for the structural images; for details, see Bendtz et al. (2022).

#### 2.4.1 Acquisition settings

All data was collected at SUBIC with a Siemens 3T Magneton Prisma MRI-scanner with a 20-channel head coil. The functional volumes were acquired with a repetition time (TR) of 2.1 s and echo time (TE) of 30 ms, with 70 slices each 2 mm thick and a 1 mm slice gap. Voxel size was 2.2 x 2.2 x 2.2 mm^3^ with a field of view (FOV) of 210 mm, and the flip angle was set to 70 degrees. Acquisition of the structural T1-weighted images was shortened for the adolescents and thus used different settings than Bendtz et al. (2022). The adolescents’ structural images were acquired with a TR of 2.3 s and TE of 2.88 ms, with 176 slices each 1.2 mm thick and a FOV of 239 mm. The duration was 2 m 32 s and was acquired directly after the experimental paradigm.

#### 2.4.2 Preprocessing, quality control, volume repair, and additional data exclusion

Preprocessing was performed with SPM12 (Friston et al., 2007) in MATLAB (The MathWorks Inc., 2019), and included realignment, coregistration of the functional images to the structural image, normalization, and spatial smoothing. Normalization utilized affine regularization, and all voxels were resampled to 2 x 2 x 2 mm^3^ using 4^th^ degree B-spline interpolation. White and grey matter segmentation and bias correction were carried out in the normalization step. Smoothing of the functional data was performed with a 3D isotropic Gaussian smoothing kernel of full-width-at-half-maximum = 8 mm.

We used MRIQC (v24.0.2, Esteban et al., 2017) to obtain quality metrics for each participant and run (to summarize over all participants and runs, we used v22.0.6). For each metric, outlier runs were checked manually whether there were any issues with a specific run’s images. We excluded 5 runs from 3 participants based on MRIQC’s estimated framewise displacement (FD), see Supplementary Materials (§S2.1) for details.

In addition to excluding data based on the FD metrics provided by MRIQC, we used ArtRepair (Mazaika et al., 2007) to repair individual volumes. We used default settings: volumes were flagged if volume-to-volume movement was above 0.5 mm (this measurement does not entirely match the algorithm used in MRIQC for FD) or if the volume-to-volume change in signal intensity exceeded 1.5%. The flagged volumes were repaired by interpolation of surrounding volumes. We excluded 5 runs from 4 subjects as the number of flagged volumes in a row exceeded 7, see Supplementary Materials (§S2.1) for more details.

### 2.5 Whole-brain analysis

#### 2.5.1 First level

We modelled the BOLD signal with eight conditions and six movement regressors. The introductory context in both the indirect and direct conditions were modelled as one regressor (regressor i). Bendtz et al. (2022) treated these as two separate regressors, however we respecified the model of the adult data to match the current setup. The question and answers were modelled separately per condition (regressors ii-v). The remaining regressors were (vi) filler trials and compliance questions; (vii) response to compliance questions, and (iix) inter-trial-interval (ITI, i.e., silence).

We modelled five different contrasts: indirect answers vs. direct answers (Indirect > Direct), direct answers vs. indirect answers (Direct > Indirect), indirect answers vs. implicit baseline (Indirect > Baseline), direct answers vs. implicit baseline (Direct > Baseline), and context vs. ITI. Our main analysis used the first two contrasts, though we included the contrasts with baseline to better understand the effect of each condition in the ROI analysis (rather than only the relative difference between them). We only used the context vs. ITI as an initial sanity check, by inspecting the individual results for this contrast in a sub-sample of the participants.

#### 2.5.2 Second level whole-brain analysis

Second level analyses were conducted using the Indirect > Direct and Direct > Indirect contrasts defined at the first level. We performed the following second level tests: (i) Indirect > Direct, all adolescents, and (ii) Indirect > Direct, Young and Mid separately. These were also performed in the opposite direction, Direct > Indirect. While we did not include the Direct > Indirect contrast for the Old sample in Bendtz et al. (2022), we have since investigated this contrast and will report these results (for LS, HS, and Old, respectively). Our main tests for group effects only used the Indirect > Direct contrast and included (iii) Young > Mid and Mid > Young; (iv) Adolescents > Old and Old > Adolescents. The cluster-forming threshold was set to *P*_uncorrected_ = .005 with Family Wise Error (FWE) as a multiple comparison correction method on the cluster and peak level. We report clusters with *P*_FWE_ < .05 and test-statistics of each cluster’s peak voxel if *P*_FWE_ < .05. Clusters with *P*_FWE_ > .05 are not reported, even if its peak voxel is *P*_FWE_ < .05.

### 2.6 ROI analyses

The ROI analyses were preregistered to address our questions regarding age-related developmental differences (RQ2—3) as well as whether differences in adult pragmatic skill may reflect delayed development (RQ4). Accordingly, two statistical approaches were used: one first testing for overall group differences followed by pairwise testing, and the second following a stepwise procedure testing for both differences and similarities between groups.

#### 2.6.1 Testing for differences with age group in literature-based ROIs

To test our hypotheses on developmental changes across age groups, we specified three ROIs: the (i) anterior medial superior frontal gyrus (amSFG); (ii) posterior medial superior frontal gyrus (pmSFG); and (iii) the left anterior temporal lobe (ATL). These ROIs are shown in Figure S5, 3, and S6, respectively.

The two ROIs in the mSFG were derived from clusters identified in the Indirect > Direct contrasts from adolescents in Asaridou et al. (2019) (amSFG) and adults in Bendtz et al. (2022) (pmSFG), respectively. As these two clusters overlap, we edited the posterior ROI by removing all voxels anterior to the most posterior voxel in the anterior ROI (MNI coordinate y = 28) using FSLeyes (Jenkinson et al., 2012). The ATL ROI was similarly derived from Asaridou et al. (2019). We manually edited the ROI to only cover anterior temporal regions, based on the definition of the ATL from Rice et al. (2015) and further restricted it to voxels present in all participants (see Supplementary Materials (§S2.4) for more details).

We extracted individual beta values (here, contrast estimates for the Indirect > Direct contrast) from each ROI using Marsbar (Brett et al., 2006) in SPM12. Subsequent analyses followed a preregistered hierarchical procedure. First, for each ROI we tested for overall group effects across the Young, Mid, and Old groups using separate ANOVAs. Second, for ROIs showing a significant effect of group (*P* < .05), we conducted preregistered pairwise T-tests (Young vs. Mid; Mid vs. Old), with correction for two tests within each follow-up. The direction of these tests was based on pre-specified hypotheses, see Table S1. This two-step approach was chosen to minimize the number of planned *T*-tests. All analyses were conducted in RStudio (Posit Team, 2024).

#### 2.6.2 Testing for potential developmental delay in the LS group

To investigate whether pragmatic difficulties in the LS group could reflect a developmental delay, we conducted a preregistered stepwise procedure. In these tests, we combined the Young and Mid into one adolescent group, as we were interested in potential similarities between the LS and adolescents. First, we identified brain regions showing differences between the HS group and adolescents in the Indirect > Direct contrast. We performed whole-brain tests in both directions (HS > Adolescents and Adolescents > HS). Significant clusters from these contrasts were used as ROIs for follow-up testing.

In a second step, we used Marsbar (Brett et al., 2006) to extract beta values (contrast estimates) for all participants from the ROIs that came out of the first step. We then tested for significant differences between extracted betas for the HS and LS groups, using T-tests. These were performed in the same direction as the tests where the clusters were revealed between the HS and adolescents (HS > Adolescents or Adolescents > HS), with the LS group replacing the Adolescents.

In a final step, we compared the beta values of the LS group and adolescents using both frequentist and Bayesian *T*-tests. The direction of these tests again matched the original test where the clusters were found, with the LS group now replacing the HS group. If the LS group shared similar activation patterns with the adolescents, we would expect non-significant results. To quantify a level of support of (no) difference, we also included Bayesian *T*-tests. All *T*-tests were performed in RStudio; for the Bayesian tests we used the BayesFactor library (Morey et al., 2024) with a default prior of √2/2.

#### 2.6.3 Extraction of beta values for the Indirect and Direct conditions

To better understand the direction of our results, we also extracted individual beta values using the Indirect > Baseline and Direct > Baseline contrast. This was done for the literature-based ROIs, any clusters yielded by group comparisons, and the ParPrec clusters from Bendtz et al. (2022). As mentioned in section 2.5.1, we only used these contrasts for descriptive purposes: consequently, we did not perform any statistical testing on this data.

#### 2.6.4 Functional cluster definition using network parcellations

To further characterize the functional profile of the clusters included in our analyses, we quantified their overlap with large-scale network parcellations. For each cluster, we counted the number of voxels overlapping with networks from one of the atlases provided by Du et al. (2024) (DU15NET-Agree53per.nii, 53% between-subject agreement). Overlap was calculated as the proportion of cluster voxels assigned to a given network relative to the number of cluster voxels assigned to any network within the atlas.

### 2.7 Supplementary analysis with only right-handed adolescents

We performed supplementary analyses in the tests involving only the adolescents, but with non-right-handed adolescents removed (n = 4). These were done in both directions (Indirect > Direct, Direct > Indirect) for the following tests: Adolescents, Young, Mid, Young > Mid, and Mid > Young. Handedness information was however not available for the Old group. The results from these tests do not change our main conclusions and are reported in Supplementary Materials §S3.2.

## 3. Results

### 3.1 Whole-brain analyses

#### 3.1.1 Indirect > Direct: All adolescents, and Young and Mid groups separately

The Indirect > Direct speech act contrast revealed several significant clusters across the brain, both when analyzing the subgroups separately (Figure 1) and when pooling all adolescents together (Figure S2A). Significant clusters and statistics are reported in Table S2.

**Figure 1.**
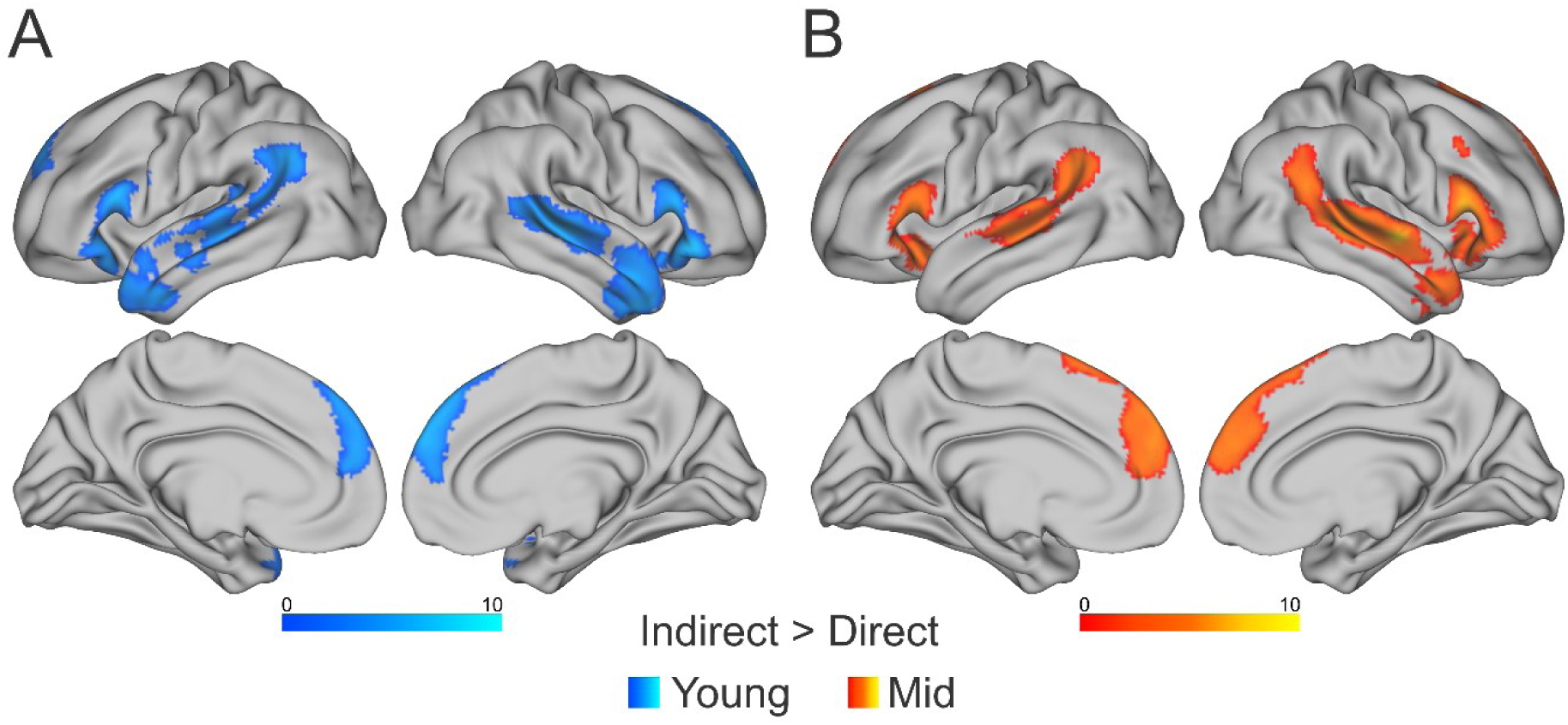
Significant clusters in the Indirect > Direct contrast for the (A) Young (ages 13-15) and (B) Mid (ages 16-18) groups separately. The figure shows clusters with a cluster-forming threshold of *P*_uncorrected_ < .005. We used family wise error (FWE) as a multiple comparison correction method on cluster and peak levels and only report clusters with a *P*_FWE_ < .05. The color scale represents *T*-values per voxel and uses the full range of *T*-values from all tests (including those in the Supplementary Materials). Significant clusters for all adolescents pooled together is provided in Figure S2A.

It is visible in the results from the two subgroups in Figure 1 that there was significant activity in both respective maps in the bilateral inferior frontal gyrus (IFG), anterior mSFG (or anterior medial prefrontal cortex), right ATL, left angular gyrus/supramarginal gyrus (AG/SMG), and the bilateral mid-to-posterior superior temporal sulcus (STS). Additionally, the Young group showed a cluster in the left ATL and the Mid group showed a cluster in the posterior mSFG (or the dorsal portion of the posterior mPFC) as well as more extensive activation in the right STS extending into the AG/SMG(see Figure 1B). The cluster in the posterior mSFG overlapped spatially with one of our planned ROIs (see Figure 3).

The overall pattern of the Indirect > Direct contrast is robust across all age groups (Young, Mid, and Old), however with important significant differences (see previous paragraph and following sections on group effects). For the Old group, the patterns are consistent with those originally reported in Bendtz et al. (2022), which is entirely expected due to using the same data (albeit with a slightly respecified model, see section 2.5.1). The results of the contrast from the Old group are presented in Figure S3, alongside the original results from Bendtz et al. (2022).

#### 3.1.2 Indirect > Direct: Group effects between the adolescent groups

No significant clusters were found when testing for group effects between the adolescent groups in either direction (Young > Mid, Mid > Young).

#### 3.1.3 Indirect > Direct: Group effects between adolescents and adults

There were no significant clusters in the Indirect > Direct contrast in the Adolescents > Old direction. The Old > Adolescents direction yielded a single significant cluster in the bilateral posterior cingulate cortex (PCC) extending into the corpus callosum (Figure S4, Table S3, spatially overlapping with the cluster in Figure 4). When we performed the same test with white matter masked out, it resulted in two smaller clusters in the same area (red outlines in Figure S4), though neither cluster reached significance (*P* = .55 and *P* = 1.00).

#### 3.1.4 Direct > Indirect: All adolescents, and Young and Mid separately

The reverse contrast (Direct > Indirect) revealed no significant clusters in the Mid group. In contrast, the Young group showed several significant clusters (Figure 2A and Table S4). This included a cluster in the left mid-to-posterior parts of the intraparietal sulcus and superior parietal lobule, located just posterior to the parietal ParPrec cluster reported in Bendtz et al. (2022). We observed further significant clusters in the bilateral anterior middle frontal gyrus (MFG), the left anterior (lateral) superior frontal gyrus (SFG), bilateral posterior ventral mPFC, and the bilateral PCC/ventral-to-mid precuneus. Pooling all adolescents together gave a similar result as with the Young group, with additional clusters in the right posterior intraparietal sulcus/SMG and left posterior SFG: these results are reported in Figure S2B and Table S4.

**Figure 2.**
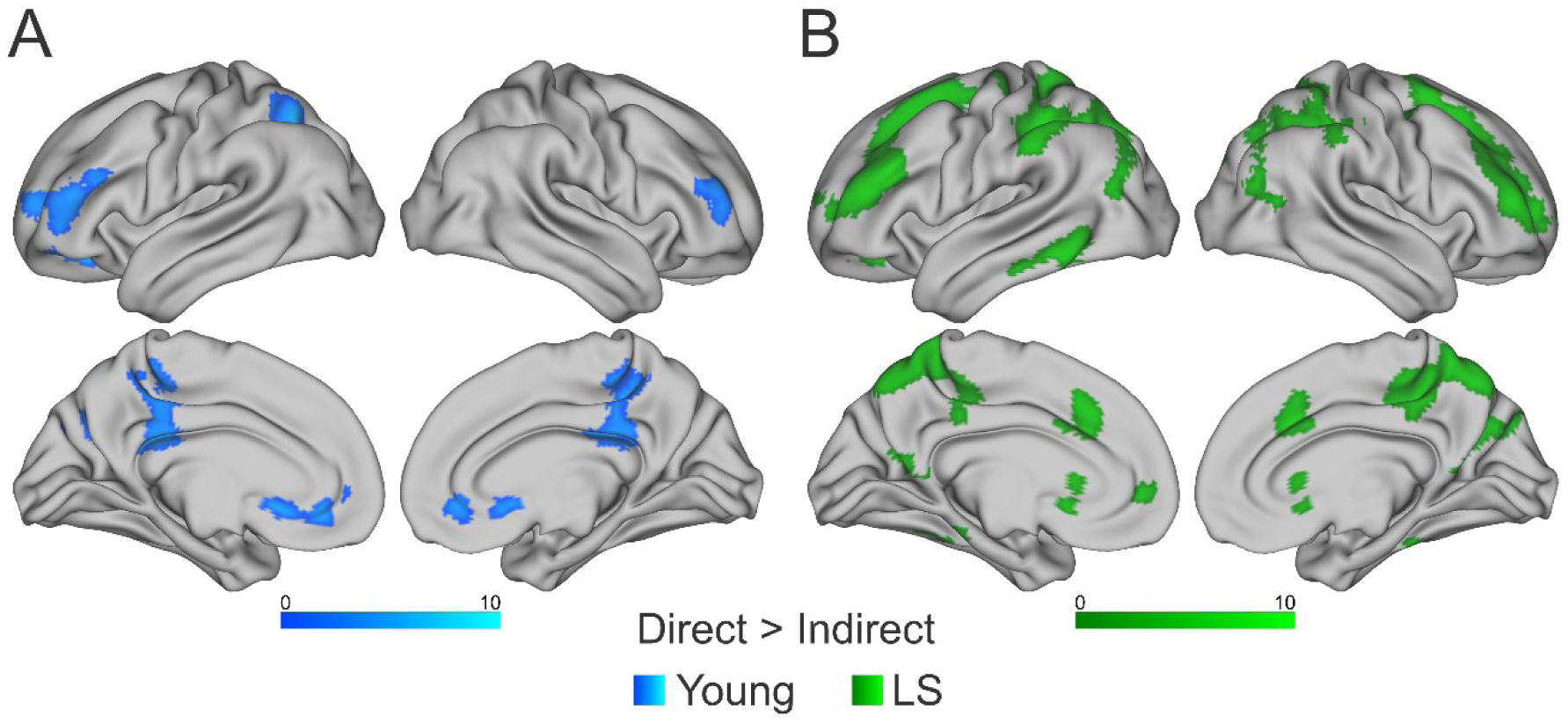
Significant (FWE corrected) clusters in the Direct > Indirect contrast for the (A) Young (ages 13-15) group and (B) LS group. There were no significant clusters in the same contrast in either the Mid or HS group. Significant clusters for all adolescents pooled together are provided in Figure S2B.

#### 3.1.5 Direct > Indirect: The Old group, and LS and HS separately

Results from the Direct > Indirect contrast in our Old sample partly resembled those in the adolescents. The contrast yielded several clusters in the LS group (Figure 2B), overlapping with regions also found in the Young group. Overlapping significant clusters were overall more extensive for the LS group (compared to the Young group), including lateral regions of the frontal lobe. Activity in the left intraparietal sulcus overlapped with the parietal ParPrec cluster, and a homologous cluster was also significant in the right hemisphere. Overlap was also found with the precuneal ParPrec cluster. Pooling all adults together rendered a similar result, however, there were no significant clusters for the HS group alone (as was also the case for the Mid group in the same contrast). The results are reported in Table S5.

### 3.2 ROI analyses

#### 3.2.1 ANOVA results and follow-up T-tests

To investigate differences across age groups (Young, Mid, and Old), we performed three ANOVAs on the Indirect > Direct contrast for ROIs in the pmSFG, amSFG, and ATL. The ANOVA for the pmSFG revealed a significant main effect of age group (*F*(2, 105) = 3.62, *P* = .03, see Figure 3). Neither of the two ANOVAs in the amSFG or the left ATL reached significance (*F*(2, 105) = 1.78, *P* = .17 (Figure S5) and *F*(2, 105) = 2.33, *P* = .10 (Figure S6)), resulting that no *T*-tests were performed for these ROIs. One of the follow-up *T*-tests in the pmSFG (one-tailed, Old > Mid) was nominally significant but did not survive Bonferroni correction (*T*(71) = -1.93, *P*_raw_ = .03). The other *T*-test (Mid > Young) did not reach significance (*T*(47) = -0.83, *P*_raw_ = .21).

**Figure 3.**
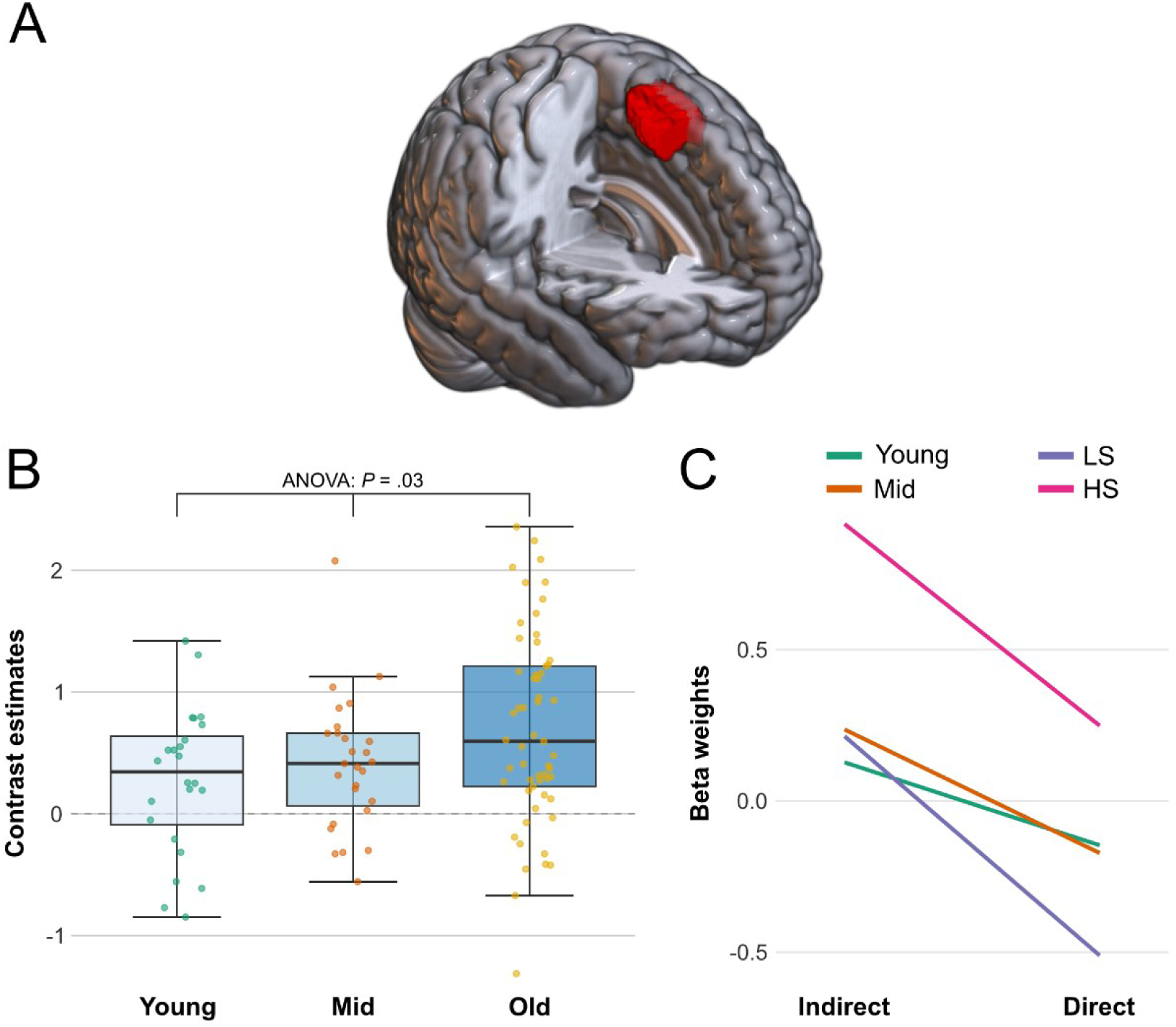
(A) Location of ROI in the posterior medial superior frontal gyrus (pmSFG). (B) Contrast estimates (Indirect > Direct) and result from ANOVA test. One of two follow-up *T*-tests (Old > Mid) was nominally significant (*P*_raw_ = .03) but did not survive Bonferroni correction. The other *T*-test (Mid > Young) was not significant (*P*_raw_ = .21). Box plots for all groups are provided in Figure S7. (C) Mean beta weights from each condition per group.

The mean contrast estimates and standard deviations for each ROI and group are reported in Table 2 and the data for the pmSFG is visualized in Figure 3 (panel B). Plots with contrast estimates for the amSFG and ATL are provided in Supplementary Materials, Figure S5-6.

**Table 2.** Mean contrast estimates for the Indirect > Direct contrast in the included ROIs.

| ROI | Young | Mid | LS | HS | Adolescents | Old |
| --- | --- | --- | --- | --- | --- | --- |
| pmSFG | 0.27 ± 0.60 | 0.41 ± 0.55 | 0.72 ± 0.72 | 0.67 ± 0.87 | 0.35 ± 0.57 | 0.70 ± 0.79 |
| amSFG | 0.79 ± 0.80 | 0.72 ± 0.74 | 1.05 ± 1.00 | 1.09 ± 0.98 | 0.75 ± 0.76 | 1.07 ± 0.98 |
| ATL | 0.36 ± 0.46 | 0.15 ± 0.36 | 0.42 ± 0.46 | 0.36 ± 0.62 | 0.25 ± 0.42 | 0.39 ± 0.54 |
| dPCC | -0.28 ± 0.48 | -0.10 ± 0.46 | -0.16 ± 0.72 | 0.47 ± 0.74 | -0.18 ± 0.47 | 0.16 ± 0.79 |

*Note: The ROIs covering the pmSFG, amSFG, and ATL were defined based on previous studies. The ROI in the dPCC was also preregistered, as all analyses in this paper, but it came from a test for group effects in the Indirect > Direct, HS > Adolescents contrast in the present study. Positive contrast estimates indicate higher activity in the Indirect condition.*

#### 3.2.2 Testing for a potential developmental delay in the LS group

To investigate our hypothesis of a potential developmental delay in the LS group, we first tested for group effects in the Indirect > Direct contrast between the HS group and adolescents (i.e. Young and Mid together). This revealed a single significant cluster in the HS > Adolescents direction, located in the dorsal aspects of the posterior cingulate cortex (dPCC) (Figure 4, Table S3). The location of this cluster was similar to the one found in the group effect Old > Adolescents (Figure S4), although more extensive in size, spanning further both posteriorly and dorsally. A test in the opposite direction (Adolescents > HS) did not result in any significant clusters.

**Figure 4.**
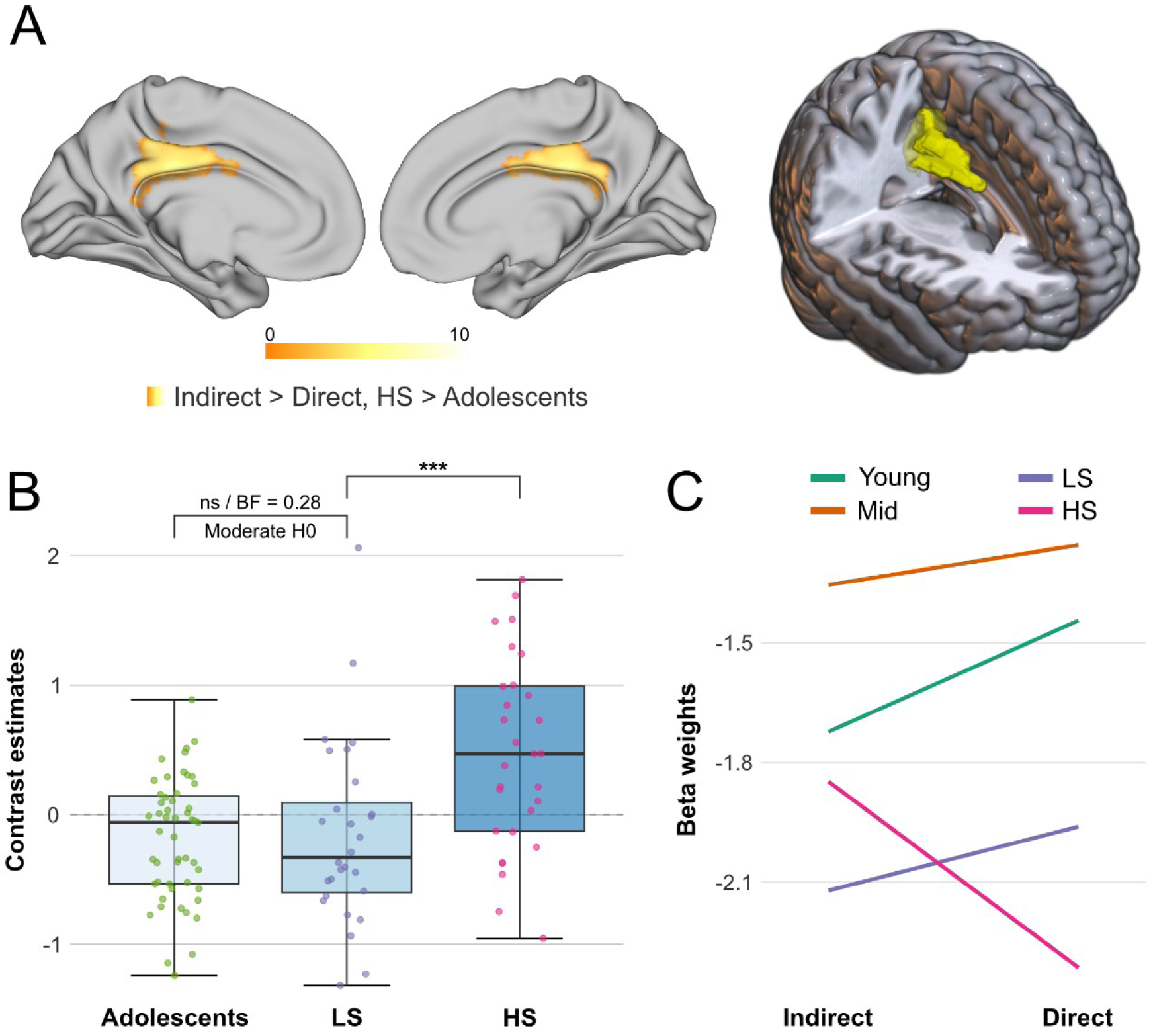
(A) Significant (with FWE-correction) cluster from whole-brain Indirect > Direct, HS > Adolescents comparison in the posterior cingulate cortex (dPCC). White matter was masked out prior to running the group test. (B) Contrast estimates (Indirect > Direct) with results from *T*-tests (Bayesian *T*-test to assess support for the null-hypothesis when comparing the Adolescents and LS groups). Star notation: *** *P*_raw_ < .001. Box plots for all groups are provided in Figure S8. (C) Mean beta weights from each condition per group.

We then extracted beta values (i.e., contrast estimates) from this cluster for all participants, separating into HS, LS, and adolescents. A one-tailed *T*-test (HS > LS) showed a significant difference between the two Old sub-groups (*T*(55) = 3.24, *P* < .001). Finally, we performed both frequentist and Bayesian *T*-tests in the direction LS > Adolescents. The frequentist *T*-test was not significant (*T*(40) = 0.16, *P* = .44) and the Bayes Factor indicated moderate support for no group difference (BF = 0.28), as hypothesized. Descriptive statistics for the contrast estimates in the dPCC, also separating adolescents into the two subgroups, are reported in Table 2. The data used in the tests is presented in Figure 4 (panel B).

#### 3.2.3 Follow-up descriptive statistics for Indirect > Baseline and Direct > Baseline

The mean beta weights of the Indirect and Direct conditions are reported in Table S6 for each ROI and group, including the dPCC cluster from the HS > Adolescents test reported in section 3.2.2. We extracted these to get a better understanding of which condition and/or group was driving the results observed in our statistical analyses. The direction of the difference between the conditions per group are plotted in each ROIs respective figure (pmSFG: Figure 3C; dPCC: Figure 4C; amSFG: Figure S5C; and ATL: Figure S6C). It should be noted that the contrast estimates reported in Table 2 and presented with boxplots in the figures reflect the relative difference between the two conditions’ beta weights (in the Indirect > Direct contrast).

Across all ROIs except the dPCC, the Indirect condition showed overall positive mean beta values. In contrast, all groups exhibited negative mean betas in the dPCC for both conditions. While negative betas here reflect relative deactivation compared to an implicit baseline, we will refer to less negative beta values in one condition relative to another as indicating higher activation, for simplicity.

In the dPCC, only the HS group showed higher activation (i.e., less negative betas) in the Indirect relative to the Direct condition. This pattern reflects a directional reversal for the HS group relative to the other groups. Mean betas of each condition in the dPCC are shown in Figure 4 (panel C). Plots for the remaining ROIs can be found in Figure 3 (pmSFG) and our supplementary materials, Figure S5-6.

As the extraction of the individual beta values for the two conditions provided additional information on the observed results in the current study, we decided to revisit the data from Bendtz et al. (2022), i.e., the ParPrec clusters, in a similar manner. We extracted betas for the Indirect > Baseline and Direct > Baseline contrasts using the ParPrec clusters as separate ROIs. The precuneal cluster resembled the pattern of the dPCC-cluster, though the mean betas were much more negative overall. In the parietal cluster, we observed that the LS group exhibited a larger difference in the direction Direct > Indirect, compared to the HS group which made little difference between conditions. The two adolescent groups also showed higher activation in the Direct relative to the Indirect condition, but less so than LS. The corresponding data are reported in Table S7 and the data is shown in Figure S9.

#### 3.2.4 Overlaps with network parcellations

Voxel-wise overlaps between the pmSFG, dPCC, and ParPrec clusters and the networks from Du et al. (2024) are provided in Table 3, only including networks with overlap >15% or where there is overlap with at least two clusters. The full set of results, including the remaining clusters (amSFG and ATL), are reported in Table S13. The pmSFG cluster mainly overlaps with LANG, and to some extent DN-B; dPCC overlaps with DN-A/B, the salience/parietal memory network (PMN) and FPN-A/B; Par overlaps with both FPN-A and the (dorsal) attention network (dATN-A); Prec overlaps with the dATN-A, PMN, and FPN-A.

**Table 3.**
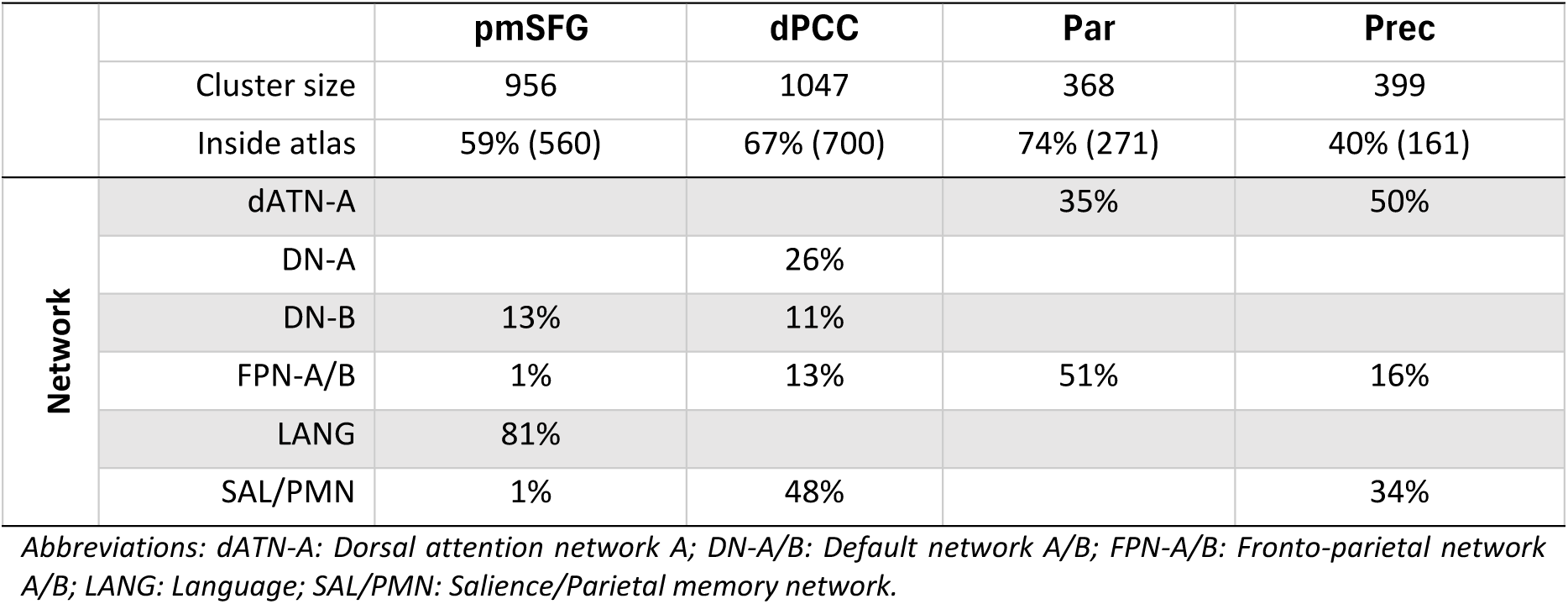
Overlaps between ROIs (pmSFG, dPCC) and the ParPrec clusters from Bendtz et al. (2022) (treated separately) and atlas from Du et al. (2024) (53% between-subject agreement). Percentage of overlapping voxels is based on the number of voxels inside atlas, rather than the original cluster.

## 4. Discussion

In the current study, we have found age-related differences in pragmatically relevant neural processes that underlie comprehension of indirect and direct speech acts. Overall, adolescents activated similar lateral and medial prefrontal areas as previously reported for adults, in an Indirect > Direct speech act contrast (Figure 1, S2-3). As hypothesized, we observed an increased recruitment across age groups (Young, 13—15 y.o.; Mid, 16—18 y.o.; Old; 20—36 y.o.) during indirect speech act (ISA) processing in the posterior medial superior frontal gyrus (pmSFG, Figure 3), which we suggest as a potential marker of pragmatic development. We found further differences in ISA activation between the high-skilled adult group (HS) and the adolescents within the bilateral dorsal posterior cingulate cortex (dPCC, Figure 4A). Consistent with our hypothesis of similarities between the low skilled adult group (LS) and the adolescents, we also gathered Bayesian support for the absence of differences in ISA activity between these groups within this cluster (Figure 4B).

As can be seen in Figure 4C, the LS, Young, and Mid groups have higher activation in the Direct versus Indirect condition in the dPCC. This pattern was stronger for the Young group compared to the Mid group. The difference between the LS and HS groups was even larger, where the HS group showed the complete opposite pattern. Hence, there is a development towards increased activity in the Indirect condition in the dPCC, that is also mirrored in the adult pattern of individual differences. This pattern is consistent with a delayed development in the LS group, as they are more similar to, in this case, both adolescent groups. We further observed a pattern of similarity between Young and LS on the one hand, and Mid and HS on the other, in the whole-brain Direct > Indirect contrast. Notably, the Young and LS groups were the only groups exhibiting significant activation in this contrast. Thus, LS participants consistently resemble the adolescents as a whole or the Young group specifically.

Our findings of age-related differences within the pmSFG (Figure 3) and dPCC (Figure 4) during pragmatic processing support the idea that pragmatics is among the abilities that continue to develop during our species’ uniquely extended childhood. This prolonged developmental period may allow for such advanced behaviors to emerge through increasing experience with diverse conversational situations. Furthermore, the results are consistent with our preregistered hypothesis that pragmatic difficulties in adulthood can reflect a delayed adolescent development. Interestingly, the two clusters that we consider to be neural markers of development and/or individual differences due to their increasing activity (again, pmSFG and dPCC) both lie outside of the perisylvian language network (although the pmSFG overlapped with the language network from Du et al., 2024), and they only partially overlapped with ToM regions. Activity within both of these networks (language and ToM) has been consistently reported in previous ISA literature (Bašnáková et al., 2014; Bendtz et al., 2022).

Taken together, these results confirm and extend on the current neuropragmatic ISA-related literature. More broadly, we contribute to the ongoing discussion within (neuro)pragmatics regarding how pragmatics relates to other cognitive domains, such as language, ToM, and cognitive control functions (Bischetti et al., 2024). Findings within the present literature (behavioral and neuropragmatic) are mixed and suggest that the relationship between domains may not only be dependent on the type of pragmatic phenomena, but also the conversational situation, developmental stage, and clinical status (Bambini et al., 2025; Frau et al., 2024; Katsos & Kissine, 2025; R. Tomasello et al., 2025). The present study adds a developmental neurobiological perspective, by demonstrating that developmental and individual differences involve additional areas beyond those usually associated with ISA.

It is noteworthy that the two clusters central to the present study (pmSFG and dPCC) both fall within third-order regions (or overlap between second-and third-order regions) when compared to a tripartite parcellation from Du et al. (2024, Figure 36). These tertiary association areas are thought to support higher-order cognitive abilities, in contrast to primary sensorimotor regions. In particular, the dPCC overlaps with the integrative, transmodal, end of the principal gradient described by Margulies et al. (2016), which encompasses the Default Mode Network (DMN, see section 4.3), as opposed to the unimodal end reflected within sensorimotor regions. Such association areas are among the last to mature in the human brain and are also disproportionally expanded relative to non-human primates (Sydnor et al., 2021; Xu et al., 2020).

It is perhaps not surprising that pragmatic comprehension and its late development, which requires complex integration of linguistic, social, and contextual information, appears to rely on these higher-order systems. From an evolutionary perspective, our observation of maturational differences in these regions contributes to the view that pragmatics may represent a uniquely developed ability in humans, part of what distinguishes us from closely related primates.

Below, we discuss the two main findings (pmSFG and the dPCC) separately, as well as reflecting on activations related to the Direct > Indirect contrast. This is followed by a broader (network) perspective on the dPCC and a final section on pragmatics’ relationship with other domains (e.g., ToM). A brief expansion on the observed activity in the pmSFG and a discussion of an additional finding in the ventromedial prefrontal cortex can be found in our Supplementary materials (§S4.1-2).

### 4.1 The posterior medial superior frontal gyrus: increasing activity across age groups

Our whole-brain analysis of the Mid group in the Indirect > Direct contrast resulted in a cluster in the posterior parts of the medial superior frontal gyrus (pmSFG), which was absent in the Young group’s results (see Figure 1). This aligns with our hypothesis of an adolescent development in the pmSFG, which was one of the prespecified ROIs for this study (Figure 3A). Indeed, the whole-brain significant cluster partially overlapped with our prespecified ROI, where we observed a significant increase for ISA activity, across the three age groups (Figure 3B).

A recent study has proposed that this area is a node in an extended language network (Wolna et al., 2025). Our overlap calculations (Table 3) indicated that our pmSFG cluster largely overlaps with language regions as defined in Du et al. (∼81%), though we want to again emphasize that it is not located within the perisylvian language network (Fedorenko et al., 2024; Hagoort, 2017; Hickok, 2009). This raises questions about what role the pmSFG may play in pragmatic processing, and why it shows increasing involvement across age groups. We have previously suggested (Forbes Schieche et al., 2025) that activity in posterior medial prefrontal areas (which the pmSFG anatomically overlaps with) can be interpreted in line with accounts of an abstract-to-context-specific gradient along the anterior– posterior axis in the frontal lobe (Badre & D’Esposito, 2009; Uddén & Bahlmann, 2012). The increased engagement of the pmSFG across age groups could indicate a developmental trajectory towards more context-specific processing and expertise—in this case driven by increased experience through repeated encounters of conversational contexts, where adolescents are highly socially motivated (e.g., peer conversations, Nippold, 1993, 2000; Raffaelli & Duckett, 1989).

### 4.2 The dorsal posterior cingulate cortex: differential roles of the default mode network in pragmatic processing

The posterior cingulate cortex (PCC) is considered a key node of the default mode network (DMN, Andrews-Hanna et al., 2010; Raichle, 2015), associated with, e.g., introspection and the construction of situation models (Menon, 2023; Ranganath & Ritchey, 2012). As shown in Figure 4C, the HS group showed greater activation within the dorsal PCC (dPCC, see Figure 5 and section 4.4 for dorsal/ventral division) in the indirect condition relative to the direct one. Both the LS group and the adolescents showed the opposite pattern. An initial interpretation of the HS group’s activation pattern may be a more efficient use of situation models, which could support rich pragmatic inferencing.

**Figure 5.**
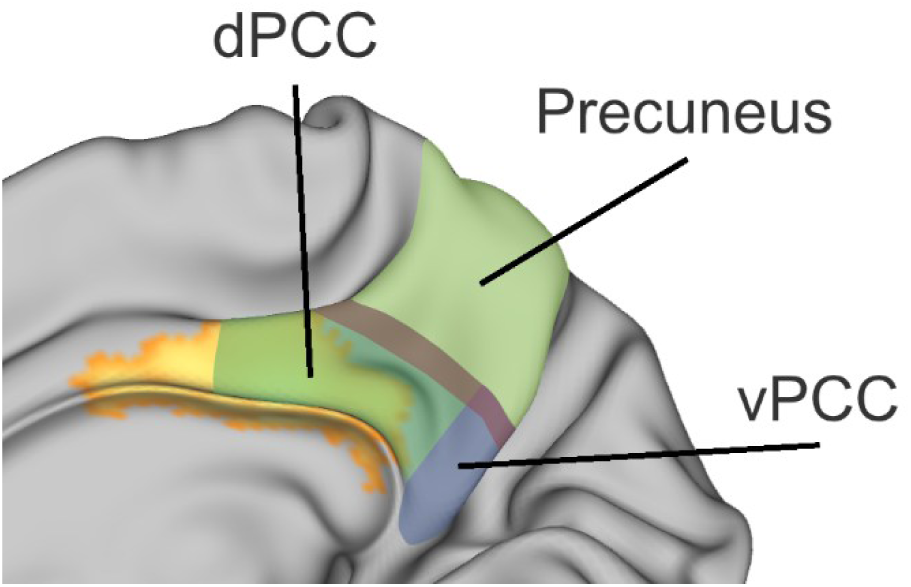
Significant cluster in the dPCC in the Indirect > Direct, HS > Adolescents contrast in yellow (medial view of the right hemisphere). Overlayed are proposed divisions of the posterior medial cortex, derived from illustrations in Foster et al. (2023) and Foster and Koslov (2025).

Activity within the PCC has hitherto not been central within other ISA studies, although e.g. Feng et al. (2021) showed increased activity within the ventral PCC related to indirect speech acts. In our data, the HS group showed significant activity in ventral regions of the PCC and precuneus in the Indirect > Direct contrast (Figure S3), whereas the LS and Young groups showed some overlapping activity with the dPCC in the reverse contrast (Figure 2). This latter activation pattern resembles results in a meta-analysis by Bohrn et al. (2012) which showed increased PCC/precuneus (though more ventrally located) activation during literal compared to figurative (idioms, irony, and metaphor) language. On the other hand, recent work in more naturalistic paradigms point to the PCC’s involvement during pragmatic and conversational processing. Thye et al. (2024) observed increased activation during narrative comprehension when the content was pragmatically richer. Extending this to more interactive settings, the PCC region has been implicated specifically during listening to conversational partners (Torubarova et al., Submitted), and has been shown to increase in involvement over the course of ongoing conversational interactions (Spatola & Chaminade, 2022). Furthermore, the PCC has also been suggested to be sensitive to differences in interpretive convergence or degree of shared interpretative mindset across individuals (Braun et al., 2025; Nguyen et al., 2019).

Despite divergent results and different perspectives in previous literature, the activity we have here observed within the dPCC appears to capture both developmental changes and individual differences in adulthood. Not only did the cluster appear in the Indirect > Direct, HS > Adolescents contrast, we also found moderate (Bayesian) support for no difference between the LS group and the adolescents. Taken together, this suggests (1) that this region may serve as a neural marker for pragmatic development and (2) that the similarity between the LS group and adolescents may indicate delayed development or extended immaturity within the LS group with respect to processing in this region during ISA. Seen alongside the pmSFG result, the observed differences suggest that pragmatic maturation is supported by multiple neural systems. We further expand on the dPCC’s role within the broader DMN and its relation to other networks in section 4.4.

### 4.3 Young and LS activity in the Direct > Indirect contrast and the ParPrec clusters

Beyond observing increased activity in the Indirect > Direct direction, we also found significant activity in the reverse contrast, Direct > Indirect for some of the groups. When separating the participants into the four subgroups (Young, Mid, LS, and HS), only the Young and LS groups exhibited significant activation (Figure 2). This increased activation may be reflective of an increased recruitment of cognitive control functions, such as working memory and cognitive flexibility. These functions are supported by the fronto-parietal Multiple Demand (MD) network (Cole, 2024; Cole & Schneider, 2007; Diachek et al., 2020), partly overlapping with the activity observed in the Direct > Indirect contrast (e.g. lateral frontal and parietal regions). Thus, the Young and LS groups may engage the MD network more when processing the direct speech acts. One possible explanation is that these groups initially consider indirect interpretations, also when processing direct speech acts. This might engage working memory processes, recruited to revisit the previous conversational context or to organize one’s thoughts to try to understand what an implicit meaning could be. The pattern of increased Direct > Indirect activity in the Young and LS group may be considered alongside our finding in the dPCC, where the Young, Mid, and LS groups showed greater activity for direct speech acts (to different extents). If increased Direct activation reflects resolving uncertainty, this may draw not only cognitive control processes supported by the MD network, but also DMN-related processes.

Furthermore, the Direct > Indirect activations (Figure 2) in the LS group was more extensive relative to the Young group. Rather than reflecting fundamentally different processes, this broader recruitment in the LS group may indicate general developmental differences in how MD (and possibly attentional) resources are engaged, consistent with findings that children and adults recruit similar MD regions, but adults show stronger activation (Schettini et al., 2023).

In addition to the overlap with the MD network, activations within the intraparietal sulcus (including the parietal ParPrec cluster in the LS group) is located near the terminations of the anterior and posterior segments of the arcuate fasciculus (Catani et al., 2005), which have been hypothesized to be associated with informative action recognition and higher level pragmatic processing (Catani and Bambini, 2014). Two further accounts may also be relevant for interpreting the involvement of the observed activations. First, a network functionally dissociated yet partially overlapping with the MD network spans both the intraparietal sulcus and the dorsal precuneus, including the ParPrec clusters (Fischer et al., 2016; Kean et al., 2025). This network has hitherto been associated with intuitive physical reasoning—i.e. making inferences about physical objects. Second, these regions also align with higher levels of a proposed cortical temporal processing hierarchy, supporting the integration of information over longer timescales (in both visual and auditory modalities, Hasson et al., 2008; Lerner et al., 2011). In our paradigm, interpreting the trial-final speech act required considering the preceding conversational context (trials being ∼18 s long), consistent with this account.

While the underlying mechanism of the Direct > Indirect activation remains to be further investigated, it is noteworthy that it was observed only in the Young and LS groups but not in the Mid or HS groups, thus reflecting differences related to both age and individual differences in adulthood.

### 4.4 Divisions within the PCC and differential roles within the larger DMN

The PCC is not a homogenous region, with prior work highlighting a subdivision into at least dorsal (dPCC) and ventral (vPCC) aspects (Foster et al., 2023; Foster & Koslov, 2025; Leech et al., 2011; Vogt et al., 2006). Our cluster from the HS > Adolescent contrast overlaps with the dPCC (see Figure 5). These subdivisions are distinguished both by cytoarchitectural features and functional properties, including connectivity patterns at rest and task responses. Functionally, the dPCC is not only a DMN node but may also act as a bridge between cognitive control networks (e.g., MD/FPN) and memory processes within the DMN and may thus be involved in the balancing of external (MD/FPN) and internal (DMN) resources (Foster et al., 2023). For our data, the HS group’s higher dPCC activity may therefore not only reflect general DMN processes, but a greater proficiency at integrating external stimuli (the verbal and voice stimuli encoding the indirect speech acts) with internal processes, such as situation models and memories of prior experiences.

A broader DMN perspective also requires considering the medial prefrontal cortex (mPFC), another core node of the DMN. The two nodes are proposed to play related but slightly different roles within the DMN (Stawarczyk et al., 2021). The mPFC node, in part overlapping with our activity in the anterior medial superior frontal gyri, has been linked to abstract, context-independent self-referential processing. By contrast, the posterior node (PCC), has been associated with context-dependent processes of both the self and others. Consistent with this distinction, all groups activated the mPFC during ISA, but only the HS group exhibited activation in the dPCC. In addition to the potential balancing role of the dPCC described above, the HS group’s strategy of pragmatic inferencing may draw not only on abstract, context-independent considerations (supported by the mPFC), but also reflect a richer, context-grounded processing mode. Notably, the context-dependent role of the posterior part of the DMN complements our findings and proposal regarding increased activity in the pmSFG (relative to the anterior mSFG). In the case of the dPCC however, both age and pragmatic skill influence its functional profile, whereas pmSFG activity differences were only related to age.

Furthermore, considering our observed activity within the DMN, we need to address the overlap between what is usually considered the ToM network and the DMN (see e.g. Neurosynth maps for “tom” and “default mode”, Yarkoni et al., 2011; Mars et al., 2012 for overlaps between the DMN and “the social brain”). Recent brain parcellations do not include a named category for ToM. Instead, the DMN is divided into subnetworks, such as DN-A and DN-B, the latter being preferentially recruited during ToM tasks (DiNicola et al., 2020; Du et al., 2024). This does not mean that the construct of ToM should be discarded. Notably, DN-A may also contribute to ToM tasks (cf. locations Du et al., 2024; Schurz et al., 2014), and, as we see in the current paper, pragmatics. Rather, ToM or mentalizing processes may be more appropriately described as subserved by subdivisions of the DMN with separate but related roles, rather than by a ToM network.

Our dPCC cluster overlaps with both the DN-A and DN-B (see Table 3), though to a greater extent with the DN-A (26% vs. 11%). It also overlaps with the MD network (∼14%, FPN-A + FPN-B), consistent with a proposed role of the dPCC as a bridge between MD and DMN (Foster et al., 2023). Notably, approximately 48% of the cluster overlaps with the salience/parietal memory network (PMN). Taken together, these overlaps reinforce the notion of the dPCC and surrounding medial parietal cortex as a convergence hub (Nijhuis et al., 2013; Weiller et al., 2025), integrating processes across DMN subdivisions, memory networks, and control systems. The HS group’s higher activity during ISA, contrasted with the opposite direction for all other groups, suggests that efficient application of these processes may serve as a marker of maturation as well as individual pragmatic differences.

### 4.5 Extending results on partial segregation between pragmatics versus language and ToM

The original results in Bendtz et al. (2022) showed two clusters where the ISA effect was greater for HS compared to LS participants: the left anterior intraparietal sulcus and the bilateral dorsal precuneus (the ParPrec clusters). These findings were taken to suggest that certain aspects of pragmatic processing can be, at least partially, segregated from structural language and ToM processes. We have since gathered further support for the pragmatic relevance of these clusters and partial segregation by examining resting-state functional connectivity in the same adult sample (Forbes Schieche et al., 2025). The ParPrec clusters showed significantly higher connectivity in the HS group, but both groups showed connectivity around zero between the ParPrec clusters and the language and ToM networks. Preliminary results from a pilot sample rather show some connectivity with the MD network, particularly for the cluster in the anterior intraparietal sulcus.

In the current study, we extend previous ISA findings in several directions. First of all, the main ISA contrast is highly similar in the adolescent and adult groups (see supplemental materials Figure S2-3), which shows the robustness of the ISA effect. Furthermore, both the described clusters (pmSFG and dPCC) were located outside the classic perisylvian language network (however, see Du et al., 2024 and Wolna et al., 2025 for the extended language network in the case of pmSFG), and rather implicating the DMN in the case of the dPCC. The locations relative to a ToM network are complex, especially in respect to the dPCC, but it is not an exclusive nor complete ToM network where we observe differences with age and/or individual differences in pragmatic skill.

Our proposed partial segregation of pragmatics and particularly ToM (Bendtz et al., 2022; Forbes Schieche et al., 2025) is part of an ongoing discussion within the broader field of pragmatics (Bambini & Lecce, 2025; Bosco et al., 2018). While neuropragmatic paradigms contrasting literal and non-literal language, such as our own contrast of ISA, often implicate ToM (see meta-analysis by Hauptman et al., 2023), both neuroimaging and behavioral studies introduce a more complex view of the involvement of ToM (Bambini & Lecce, 2025). In Tomasello et al.’s (2025) review of several neuroimaging studies, speech act type in itself (e.g., request vs. promise) differentially involves ToM, even if both are expressed directly.

### 4.7 Limitations

The present study included a new sample of adolescents to examine pragmatic development. While the main Indirect > Direct contrast was highly similar across adolescent and adult groups, supporting the robustness of the ISA effect, the study is cross-sectional, meaning that age serves only as an imperfect proxy for developmental stage.

A limitation of the present study concerns the construction of the stimuli. Although the direct and indirect replies were matched linguistically, they differ in speech act type (informative response versus criticism), and only the indirect replies serve a face-saving function. Consequently, we cannot completely disentangle these aspects in our contrast (see e.g., Boux et al., 2023). Nonetheless, our contrast is undoubtedly pragmatic in nature. While we have provided an initial insight into adolescent neural pragmatic development, it remains to be seen how these findings generalize more broadly to the full class of indirect speech acts as well as other pragmatic phenomena.

Regarding the Direct > Indirect effect observed within the Young and LS groups, we acknowledge that it may be partially amplified by the structure of the task. Still, the effect is almost certainly related to pragmatics. The adult participants were separated based on pragmatic ability outside the scanner and we observed significant activation in the Direct > Indirect contrast only among LS adults as well as the Young adolescent group. Therefore, we argue that the observed effect is pragmatically relevant— grounded in both pragmatic development and adult individual differences. A future direction is to investigate the extent to which our suggestion of overinterpretation of direct speech acts occurs in more natural settings.

## Conclusion

The outcomes of the present study provide evidence for the development of pragmatic processing from adolescence into adulthood. Neurobiologically, this development occurs outside the classic perisylvian language network, for instance (1) activity extending in a posterior direction along the medial superior frontal gyrus and (2) in the default mode network in the posterior cingulate cortex, which potentially reflects increased context specificity. Moreover, as we observed similarities between adult individuals with challenged pragmatic functioning and adolescents, we suggest that pragmatic challenges in adulthood reflect a delayed developmental path. Taken together, our results suggest that pragmatic maturation does not reflect a developmental trajectory within a single network but is rather supported by multiple networks that may mature at different rates and contribute complementary aspects of pragmatic processing.

## Acknowledgements

The authors want to thank Katarina Bendtz and Josephine Schneider for their role in the early phase of this project, e.g., collecting the behavioural and MR data of the adult sample, and Josephine Schneider for her contribution to the ethical application supporting the adolescent data acquisition. We also thank Ronja Roll, Johanna Sundström, Naira Sardarian, and Annika Andersson for their assistance in collecting the adolescent data. Data acquisition was supported by a grant to the Stockholm University Brain Imaging Centre (SU-FV-5.1.2-1035-15).

## Declaration of generative AI and AI-assisted technologies in the manuscript preparation process

During the preparation of this work the first author used ChatGPT (free version) for suggestions for improving readability and structure. The authors take full responsibility for the final manuscript.

