## Supplementary Materials for "Adolescent pragmatic development mirrors pragmatic individual differences in adulthood: An fMRI study"

### S1. Supplementary introduction

### S2. Supplementary method

#### S2.1 Excluded data and quality control

One participant withdrew participation soon after starting the first run. One run was excluded due to volume issues, another due to the participant falling asleep during the run. Some further individual runs were excluded on the basis of how many wrong answers participants gave to the control questions in a given run. Per run, we allowed for one error. 10 individual runs from 9 participants were removed.

We discovered susceptibility artifacts in two participants when inspecting individual structural and functional images after running MRIQC (Esteban et al., 2017). The artifacts were large enough to appear as warping of the brain in the structural image and signal loss in the functional images. Data from both participants was completely excluded.

Using framewise displacement (FD) as estimated by MRIQC, we excluded runs where average FD > 0.25 mm or if more than 20% of the FD values were above 0.5 mm. Following these thresholds, 5 runs from 3 participants were excluded.

Running ArtRepair (Mazaika et al., 2007) to repair individual volumes, the mean number of volumes flagged and repaired were 3.6 per run (± 6.5). 17 participants had no volumes repaired. As ArtRepair uses interpolation of surrounding volumes for repairing flagged volumes, too many flagged volumes in a row would mean that different conditions would be given the same data. We set the limit at 7 volumes in a row, equating to 14.7 s (repetition time in the scanner was 2.1 s).

Furthermore, we ran first level analysis on both the unrepaired and repaired data. We then used the art_summary function in ArtRepair to compare the two models. If repairing data led to smaller standard deviation and residual mean square in a participant’s model, we used the repaired model in the second level analysis: if either of the two metrics were not improved, we kept the original.

#### S2.2 Preregistration

Before running any analysis, but after preregistration, we decided to collect more data to replace excluded participants. Our final sample (n = 55) was therefore larger than the one described within the preregistration (n = 47). Hence, attentive readers of both the preregistration and the article may notice that the number of adolescent participants does not match between these documents.

#### S2.3 Stimulus

As mentioned in the article, the main stimulus – recorded conversations – was not changed. However, we did change some of the control questions. This was done to have them more detailed rather than general and to better balance where in the conversation the answer to the control question appeared (context, question, or answer). For example, throughout one conversation the word “car” is mentioned multiple times, and its original control question was “Was there someone speaking about cars?”. This question was changed to “Was there someone who had just bought a car?”, as this was mentioned only once in the context part of the conversation (for both its indirect and direct versions). All control questions, including those that were not changed, were re-recorded by the first author in an anechoic chamber at the Department of Linguistics at Stockholm University.

##### S2.3.1 Trial order

There were eight versions of the paradigm, divided into two sets. Within each set, the same trials were assigned to the same runs across versions. The four versions per set were organized into two pairs. Within each pair, trial order was identical, but the direct/indirect version of the trial was interchanged.

#### S2.4 ROI analysis

The ROI in the left anterior temporal lobe (ATL) was sourced from the adolescents in Asaridou et al. (2019). However, this cluster covered both anterior and posterior parts of the temporal lobe. Therefore, we decided to edit the cluster to only include more anterior parts. We used a definition of the ATL from Rice et al. (2015): we only included voxels anterior to a diagonal plane with an anterior/dorsal edge at y = 0 and z = -5, and a posterior ventral edge at y = -20 and z = 30.

| Table S1. Direction of follow-up *T*-tests. Note that the *T*-tests were only performed if the ROIs’ respective ANOVA gave a significant result. < and > indicates one-sided test, ≠ indicates two-sided. | | | | | |
| --- | --- | --- | --- | --- | --- |
| Increased activity with age | | | | | |
| pmSFG | Young | < | Mid | < | Old |
| Left ATL | Young | < | Mid | < | Old |
| Decreased activity with age (at least between adolescence and adulthood) | | | | | |
| amSFG | Young | ≠ | Mid | > | Old |

#### S2.5 Supplementary analysis using non-default settings

The main analysis, as reported in the article, used default masking settings in SPM during first- and second-level analysis. In addition to this, we ran a supplementary analysis using non-default masking procedures. This decision was based on an observation—after the whole-brain second-level analysis was run—that voxel data was missing in certain regions, primarily in the anterior frontal and inferior temporal lobes. Some missing data can be expected in these areas, as signal loss (or reduction) is common in these locations (Rua et al., 2018). We carried out the supplementary analysis to assess whether we could observe any meaningful differences compared to the main analysis. Results from this analysis are reported only in the supplementary materials (see §3.1). For reference, heatmaps showing which voxels had missing data in some participants are provided in Figure S1. The following paragraphs describe the steps of the non-default procedure.

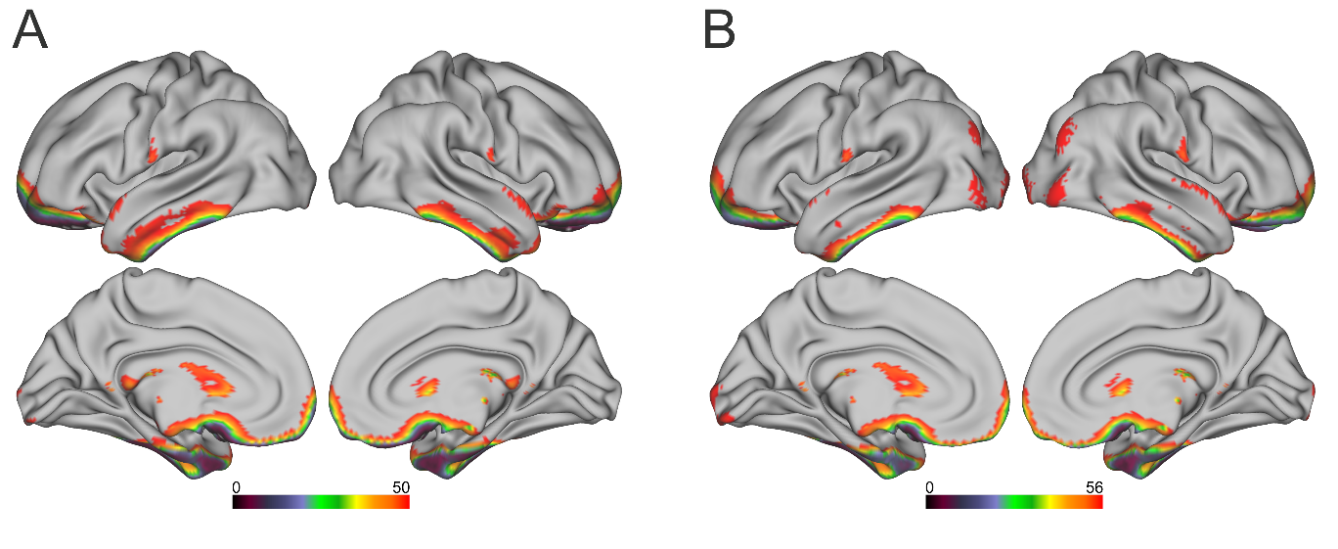

**Figure S1.** Heatmaps over the number of participants with valid (non-NaN) data in voxels for (A) adolescents (n = 51) and (B) adults (n = 57). Grey areas indicate where all participants had valid data. The scales’ respective upper limits represent where one participant had invalid data in a particular voxel (not necessarily the same participant in all locations).

On the first level, we lowered the default masking threshold in SPM from 0.8 to 0.6 for the adolescent participants. This allowed more voxels to be included per participant, even voxels with lower signal (i.e., voxels with an estimated signal strength of at least 60% of the overall mean were included).

By default, SPM’s second-level analysis applies an implicit mask based on all included participants. This mask excludes voxels missing in even a single participant. To address this, we performed the following steps: (i) We combined the first-level masks from each participant and created a binary mask in which voxels were included if they contained valid data from at least one participant. (ii) We replaced NaN values with zeros (representing no difference between conditions) in each participant’s contrast images for Indirect > Direct and Direct > Indirect. These NaN values resulted either from signal loss or when participants moved out of the initially set field of view (FOV). (iii) We used the combined mask as an explicit mask at the second level, using the modified contrast images. This explicit masking had two effects: (a) it reduced the number of non-brain voxels that had been assigned a non-NaN value during the replacement step; (b) it allowed for maximum brain coverage, as it included voxels with valid data from even a single participant. Although this approach involved adding dummy data (i.e., zeros replacing NaN values), we argue that it does not inflate the risk of false positives, as the inserted values represent no difference between conditions. In any voxel with added zeros in some participants, the original valid data must show stronger effect sizes to obtain a significant result, as the addition of zero values shifts a voxel’s estimate toward no effect. However, we note that any differences in significant results compared with those from the default procedure should be interpreted with care.

### S3. Supplementary results

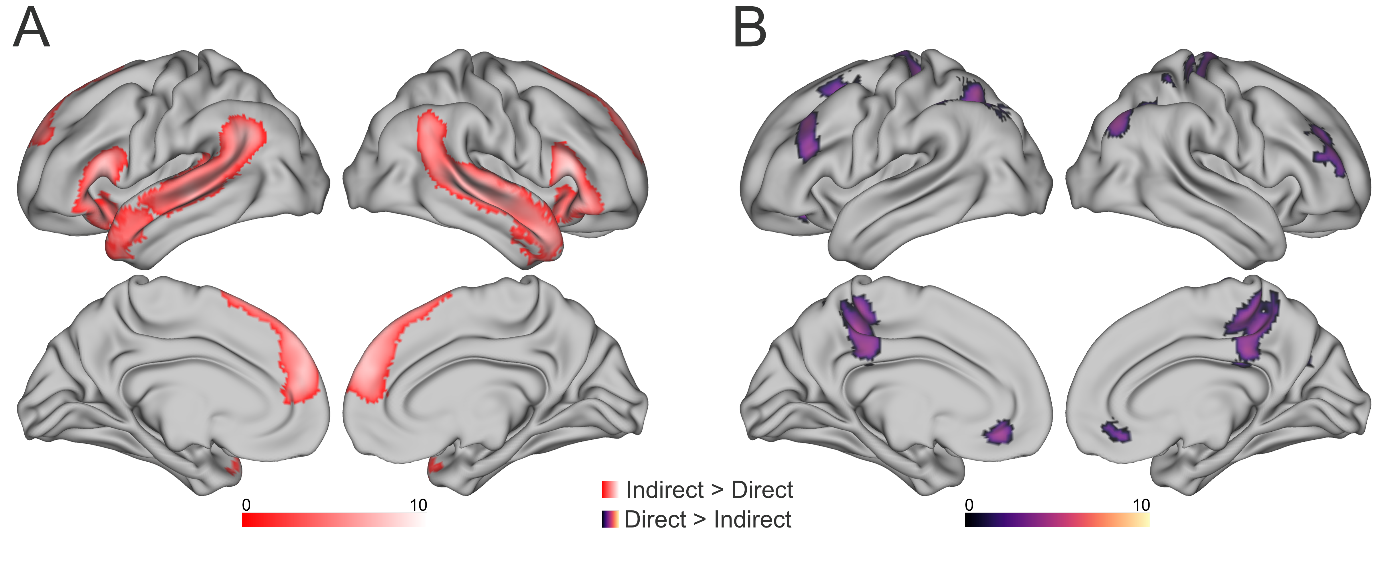

**Figure S2**. Significant clusters in the (A) Indirect > Direct and (B) Direct > Indirect contrasts, with the Young and Mid group pooled together.

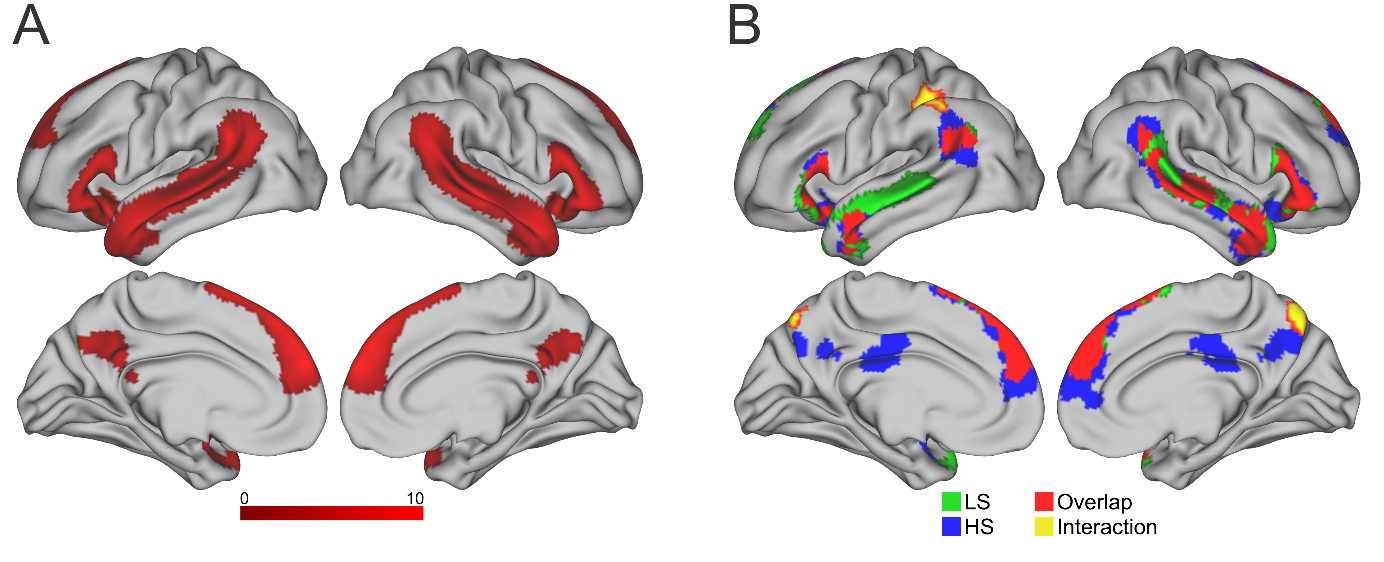

**Figure S3.** Significant clusters from the Indirect > Direct contrast in the Old group. Panel A shows all adults pooled together, using the respecified model for this paper. The respecified model treated context as one condition rather than two (i.e., regardless if it preceded an indirect or direct reply) as was originally done in Bendtz et al. (2022). Panel B shows the original results from Bendtz et al. (2022), separated into the two groups HS and LS, as well as where the two overlap in significant activation. Yellow shows the ParPrec clusters which were revealed in an interaction between the two groups (HS > LS). Note that Panel A used a cluster-forming threshold of *P*_uncorrected_ = .005 to match the adolescent threshold in this paper, while Bendtz et al. (2022) (Panel B) used a slightly more conservative threshold, *P*_uncorrected_ = .001. Also note that activations within the posterior cingulate cortex in the Old group (Panel A, all adults together) were only present for the HS group in the results from Bendtz et al. (2022).

| Table S2. Significant clusters from the Indirect > Direct contrast for all adolescents and the Young and Mid group separately | | | | | | | |
| --- | --- | --- | --- | --- | --- | --- | --- |
| Anatomical region | **Local maxima (MNI)** | | | **Cluster** | | **Voxel** | |
|  | x | y | z | k | *P*_FWE_ | *T*-value | *P*_FWE_ |
| All adolescents |  |  |  |  |  | *T*(50) |  |
| Right M/STG/ATL/IFG/AG | 58 | -34 | 2 | 6080 | < .001 | 8.76 | < .001 |
| Bilateral mSFG/mPFC/anterior cingulate | -8 | 54 | 28 | 3948 | < .001 | 7.84 | < .001 |
| Left M/STG/ATL/IFG/AG/SMG | -54 | 20 | 10 | 5337 | < .001 | 7.47 | < .001 |
| Left cerebellum Crus I/II | -20 | -80 | -34 | 505 | .016 | 5.87 | .011 |
| Young > Mid | | | | | | | |
| No significant clusters | | | | | | | |
| Mid > Young | | | | | | | |
| No significant clusters | | | | | | | |
| Young | | | | | | *T*(23) |  |
| Right IFG/ATL | 50 | 32 | -8 | 1931 | < .001 | 7.25 | .013 |
| Bilateral anterior mSFG/mPFC | 8 | 58 | 26 | 2160 | < .001 | 7.01 | .020 |
| Right M/STG | 56 | -34 | -2 | 1685 | < .001 | n.s. | |
| Left IFG/ATL | -60 | 18 | 18 | 1502 | < .001 | n.s. | |
| Left posterior STG/AG/SMG | -50 | -58 | 32 | 1975 | < .001 | n.s. | |
| Mid | | | | | | *T*(26) |  |
| Right M/STG/AG/ATL | 62 | -10 | -4 | 2711 | < .001 | 8.76 | < .001 |
| Left IFG/insula | -24 | 20 | -12 | 933 | < .001 | 7.07 | .010 |
| Right IFG/insula | 52 | 22 | 10 | 1309 | < .001 | 6.92 | .013 |
| Left M/STG/AG/SMG | -52 | -26 | -2 | 1624 | < .001 | 6.43 | .037 |
| Bilateral anterior/posterior mSFG/mPFC | -8 | 52 | 26 | 2721 | < .001 | n.s. | |
| Left cerebellum crus I/II | -20 | -76 | -28 | 617 | .003 | n.s. | |

| Table S3. Clusters from tests for group effects in the Indirect > Direct contrast, Adolescent > Old and Old > Adolescents, Adolescents > HS, and HS > Adolescents. When removing white matter, the significant cluster Old > Adolescents did not reach significance and was separated into two clusters. | | | | | | | |
| --- | --- | --- | --- | --- | --- | --- | --- |
| Anatomical region | **Local maxima (MNI)** | | | **Cluster** | | **Voxel** | |
|  | x | y | z | k | *P*_FWE_ | *T*-value | *P*_FWE_ |
| Adolescents > Old | | | | | | *T*(106) |  |
| No significant clusters | | | | | | | |
| Old > Adolescents | | | | | | *T*(106) |  |
| Dorsal posterior cingulate cortex | 4 | -28 | 22 | 575 | .010 | n.s. | |
| Old > Adolescents (white matter removed) | | | | | | *T*(106) |  |
| Dorsal posterior cingulate cortex | 8 | -26 | 28 | 230 | n.s. | n.s | |
| Dorsal posterior cingulate cortex | -2 | 44 | 36 | 64 | n.s. | n.s. | |
| Adolescents > HS | | | | | | *T*(78) |  |
| No significant clusters | | | | | | | |
| HS > Adolescents | | | | | | *T*(78) |  |
| Dorsal posterior cingulate cortex | 0 | -28 | 26 | 1047 | .003 | 5.71 | .003 |

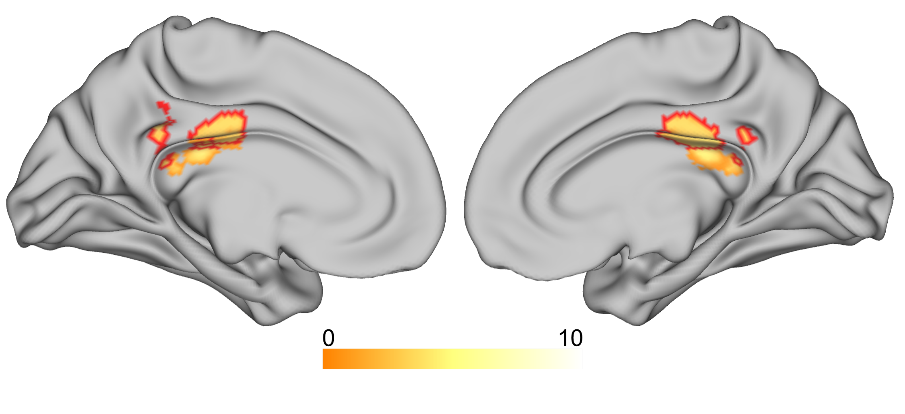

**Figure S4.** Clusters from the Indirect > Direct, Old > Adolescents comparison in the posterior cingulate cortex (PCC). Orange/yellow cluster was significant (*P* = .010) and extended into white matter in the corpus callosum. The red outline shows results from the same comparison with white matter removed. This resulted in two separate clusters, neither of which reached significance. Detailed results from both tests are reported in Table S3.

| Table S4. Significant clusters from the Direct > Indirect contrast for all adolescents and the Young and Mid group separately. | | | | | | | |
| --- | --- | --- | --- | --- | --- | --- | --- |
| Anatomical region | **Local maxima (MNI)** | | | **Cluster** | | **Voxel** | |
|  | x | y | z | k | *P*_FWE_ | *T*-value | *P*_FWE_ |
| All adolescents |  |  |  |  |  | *T*(50) |  |
| Bilateral dPCC/precuneus/left anterior intraparietal sulcus | -26 | -24 | 30 | 3935 | < .001 | n.s. | |
| Left mid-to-posterior intraparietal sulcus/superior parietal lobule/middle occipital gyrus | -30 | -68 | 54 | 1000 | < .001 | n.s. | |
| Left posterior M/SFG | -28 | 22 | 60 | 456 | .027 | n.s. | |
| Medial-to-left orbitofrontal cortex | -20 | 34 | -14 | 528 | .013 | n.s. | |
| Right AG/posterior intraparietal sulcus/middle occipital gyrus | 40 | -72 | 40 | 461 | .026 | n.s. | |
| Left MFG/IFG (triangular part) | -36 | 32 | 36 | 658 | .004 | n.s. | |
| Right anterior MFG | 34 | 40 | 46 | 776 | .001 | n.s. | |
| Right precentral and postcentral gyrus | 18 | -20 | 66 | 448 | .029 | n.s. | |
| Young |  |  |  |  |  | *T*(23) |  |
| Left M/SFG/IFG (triangular part)/orbitofrontal cortex | -20 | 30 | -12 | 1952 | < .001 | n.s. | |
| Left mid-to-posterior intraparietal sulcus/precuneus/cuneus | -32 | -62 | 54 | 1030 | < .001 | n.s. | |
| Bilateral dPCC/PCC/precuneus | 18 | -44 | 48 | 1489 | < .001 | n.s. | |
| Right anterior MFG | 44 | 56 | 12 | 513 | .013 | n.s. | |
| Mid |  |  |  |  |  |  |  |
| No significant clusters | | | | | | | |

| Table S5. Significant clusters from the Direct > Indirect contrast for all adults and the HS and LS groups separately. | | | | | | | |
| --- | --- | --- | --- | --- | --- | --- | --- |
| Anatomical region | **Local maxima (MNI)** | | | **Cluster** | | **Voxel** | |
|  | x | y | z | k | *P*_FWE_ | *T*-value | *P*_FWE_ |
| All adults |  |  |  |  |  | *T*(56) |  |
| Left middle and superior frontal gyrus/bilateral precuneus/intraparietal sulcus/superior parietal lobe | -28 | -66 | 40 | 10971 | < .001 | 6.48 | .001 |
| Left posterior inferior temporal gyrus/fusiform gyrus | -54 | -52 | -14 | 1277 | < .001 | n.s. | |
| Right M/SFG | 44 | 46 | 18 | 3241 | < .001 | n.s. | |
| Right cuneus/occipital superior gyrus | 26 | -58 | 20 | 478 | .024 | n.s. | |
| LS |  |  |  |  |  | *T*(27) |  |
| Bilateral dPCC/vPCC/precuneus/intraparietal sulcus | -42 | -46 | -46 | 8223 | < .001 | 7.88 | .001 |
| Left posterior inferior temporal gyrus/Cerebellum Crus I/II/VI/VIII | -40 | -42 | -30 | 1893 | < .001 | 7.46 | .003 |
| Left middle and superior frontal gyrus | -24 | 40 | -10 | 4454 | < .001 | 7.09 | .008 |
| Right middle and superior frontal gyrus | 32 | 56 | 8 | 3121 | < .001 | n.s. | |
| Right Cerebellum crus VIII/VI/I | 28 | -36 | -38 | 445 | .019 | n.s. | |
| Right Cerebellum Crus II/I/VIII | 44 | -66 | -44 | 475 | .013 | n.s. | |
| Bilateral middle and anterior cingulate cortex | 10 | 22 | 32 | 503 | .010 | n.s. | |
| Right ventral PCC/cuneus | 14 | -52 | 20 | 447 | .018 | n.s. | |
| HS |  |  |  |  |  |  |  |
| No significant clusters | | | | | | | |

| Table S6. Mean beta weights for the Indirect and Direct conditions separately. | | | | | | |
| --- | --- | --- | --- | --- | --- | --- |
| ROI | **Young** | **Mid** | **LS** | **HS** | **Adolescents** | **Old** |
| Indirect condition (Indirect > Baseline) | | | | | | |
| pmSFG | 0.13 ± 0.81 | 0.24 ± 1.33 | 0.21 ± 1.68 | 0.92 ± 1.36 | 0.19 ± 1.11 | 0.57 ± 1.55 |
| amSFG | 0.87 ± 1.31 | 0.67 ± 2.21 | 0.63 ± 1.96 | 1.01 ± 1.70 | 0.76 ± 1.82 | 0.83 ± 1.82 |
| ATL | 2.54 ± 0.93 | 2.33 ± 1.18 | 3.30 ± 1.39 | 3.57 ± 1.31 | 2.43 ± 1.06 | 3.44 ± 1.34 |
| dPCC | -1.72 ± 0.95 | -1.35 ± 0.96 | -2.12 ± 0.91 | -1.85 ± 1.15 | -1.53 ± 0.96 | -1.98 ± 1.04 |
| Direct Condition (Direct > Baseline) | | | | | | |
| pmSFG | -0.15 ± 0.88 | -0.17 ± 1.24 | -0.51 ± 1.64 | 0.25 ± 1.30 | -0.16 ± 1.07 | -0.12 ± 1.51 |
| amSFG | 0.07 ± 1.16 | -0.05 ± 1.81 | -0.42 ± 1.63 | -0.08 ± 1.69 | 0.01 ± 1.52 | -0.25 ± 1.66 |
| ATL | 2.18 ± 0.92 | 2.20 ± 1.24 | 2.87 ± 1.51 | 3.21 ± 1.37 | 2.19 ± 1.09 | 3.04 ± 1.44 |
| dPCC | -1.44 ± 0.92 | -1.25 ± 0.87 | -1.96 ± 1.03 | -2.31 ± 1.30 | -1.34 ± 0.89 | -2.14 ± 1.18 |
| Table S7. Mean beta weights for the Indirect and Direct conditions separately in the parietal and precuneal clusters from Bendtz et al. (2022). | | | | | | |
| ROI | **Young** | **Mid** | **LS** | **HS** | **Adolescents** | **Old** |
| Indirect condition (Indirect > Baseline) | | | | | | |
| Par | -1.80 ± 1.29 | -1.23 ± 1.39 | -2.41 ± 1.50 | -1.41 ± 1.23 | -1.49 ± 1.36 | -1.90 ± 1.45 |
| Prec | -5.34 ± 3.05 | -5.18 ± 3.09 | -5.98 ± 2.72 | -4.65 ± 2.44 | -5.26 ± 3.04 | -5.30 ± 2.64 |
| Direct Condition (Direct > Baseline) | | | | | | |
| Par | -1.49 ± 1.25 | -1.05 ± 1.25 | -1.65 ± 1.44 | -1.44 ± 1.24 | -1.26 ± 1.26 | -1.54 ± 1.34 |
| Prec | -5.03 ± 2.77 | -5.04 ± 2.64 | -4.88 ± 2.71 | -5.09 ± 2.73 | -5.03 ± 2.67 | -4.99 ± 2.70 |

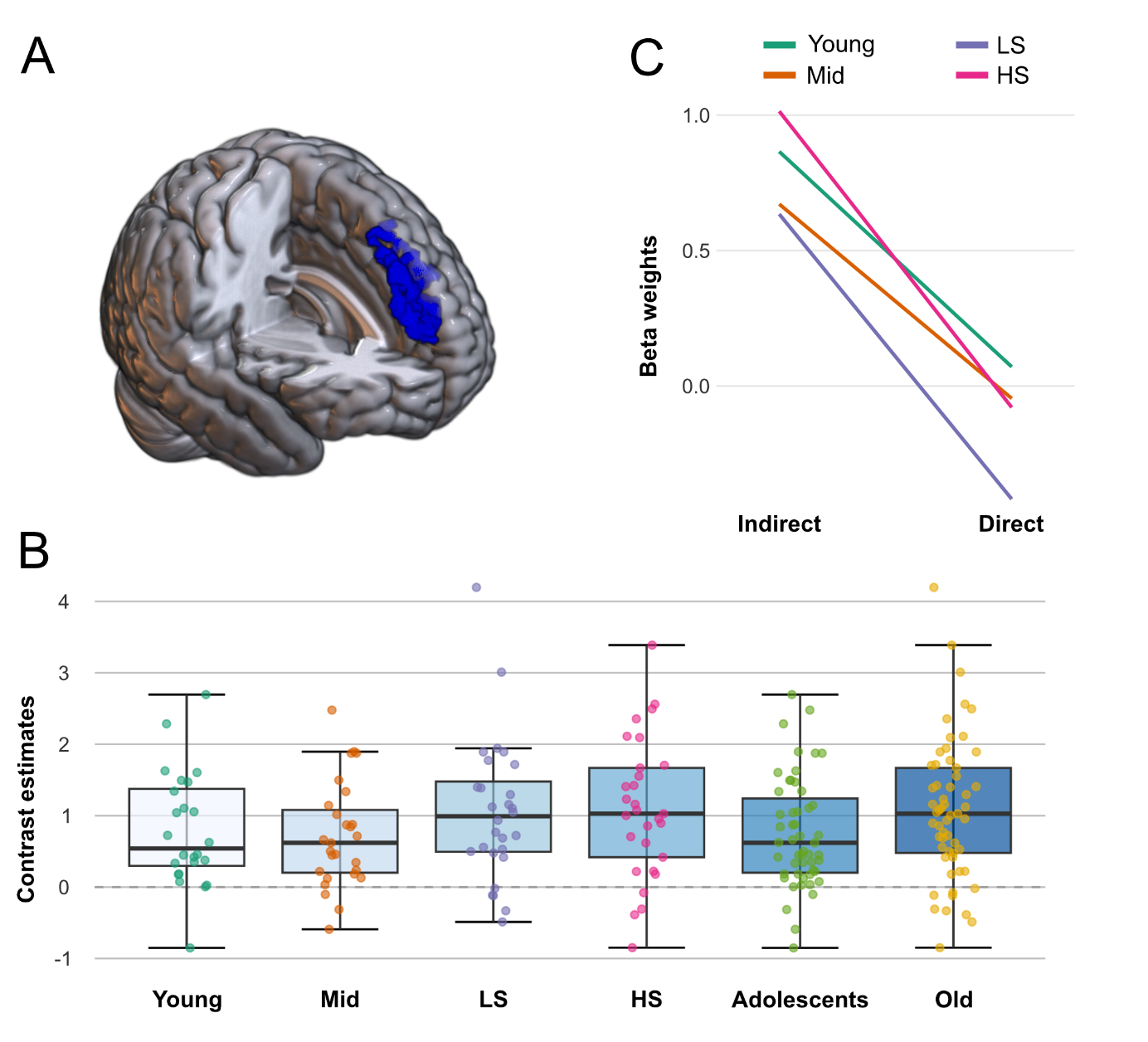

**Figure S5.** (A) Location of a ROI in the anterior medial superior frontal gyrus (amSFG). (B) Contrast estimates (Indirect > Direct) for each group, including the pooled adolescent and adult (Old) groups. An ANOVA test including the Young, Mid, and Old groups did not reach significance (*P* = .14). (C) Mean beta weights from each condition per group.

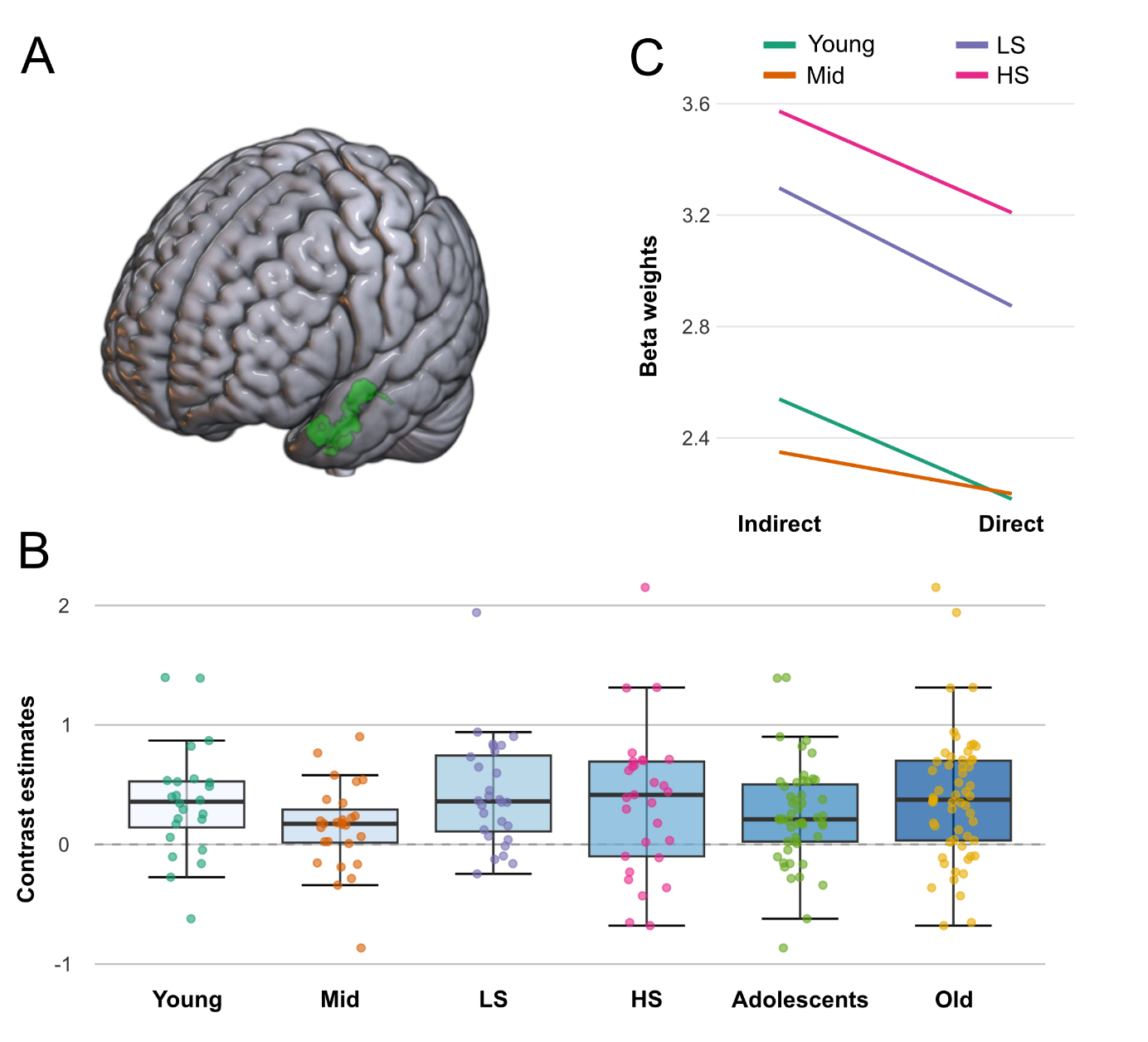

**Figure S6.** (A) Location of a ROI in the left anterior temporal lobe (ATL). (B) Contrast estimates (Indirect > Direct) for each group, including the pooled adolescent and adult (Old) groups. An ANOVA test including the Young, Mid, and Old groups did not reach significance (*P* = .10). (C) Mean beta weights from each condition per group.

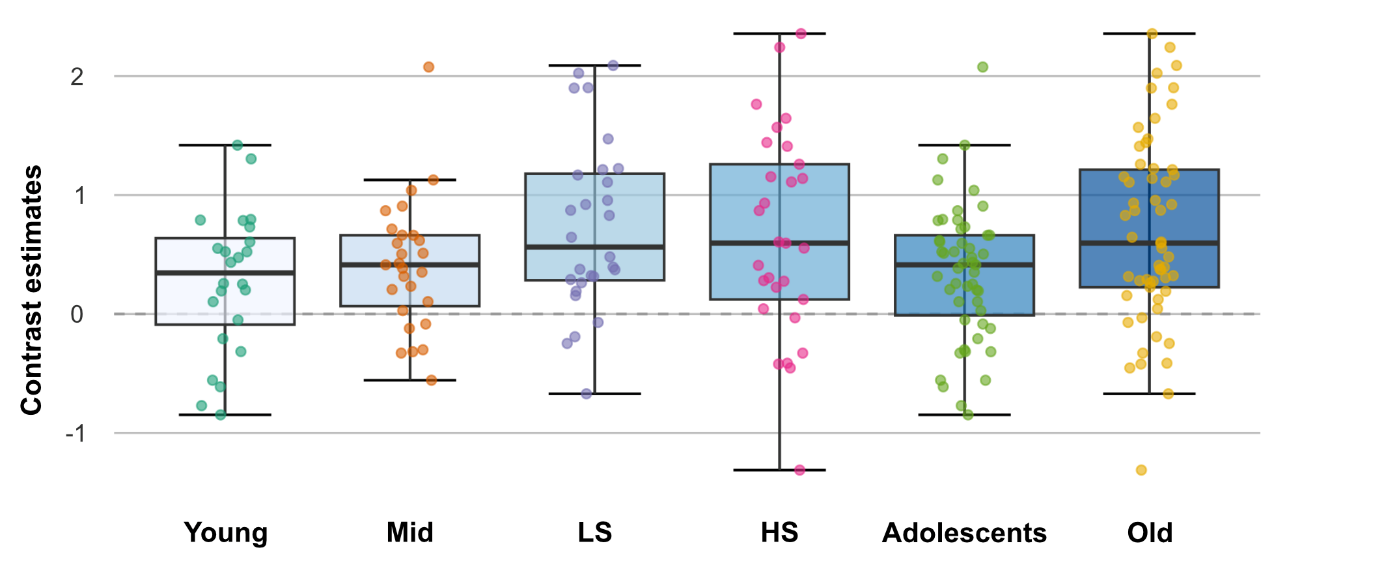

**Figure S7.** Contrast estimates (Indirect > Direct) for each group in the ROI in the posterior medial superior frontal gyrus (pmSFG). For location of this ROI, see Figure 3 in the main manuscript.

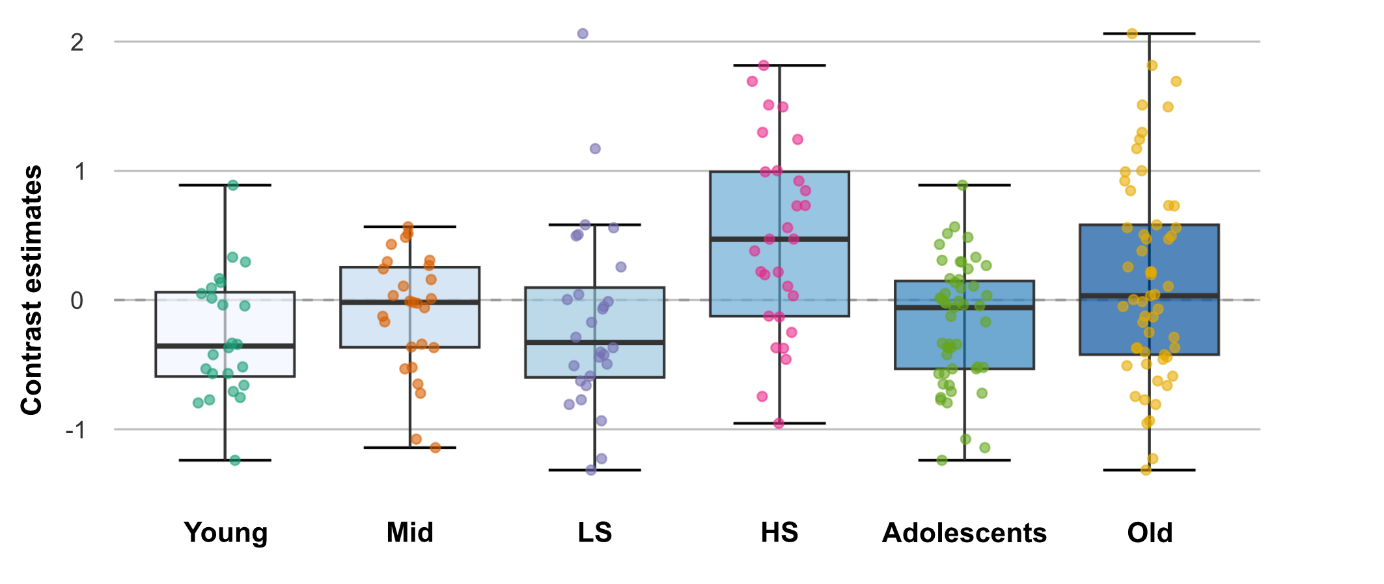

**Figure S8.** Contrast estimates (Indirect > Direct) for each group in the ROI in the dorsal posterior cingulate cortex (dPCC). For location of this ROI, see Figure 5 in the main manuscript.

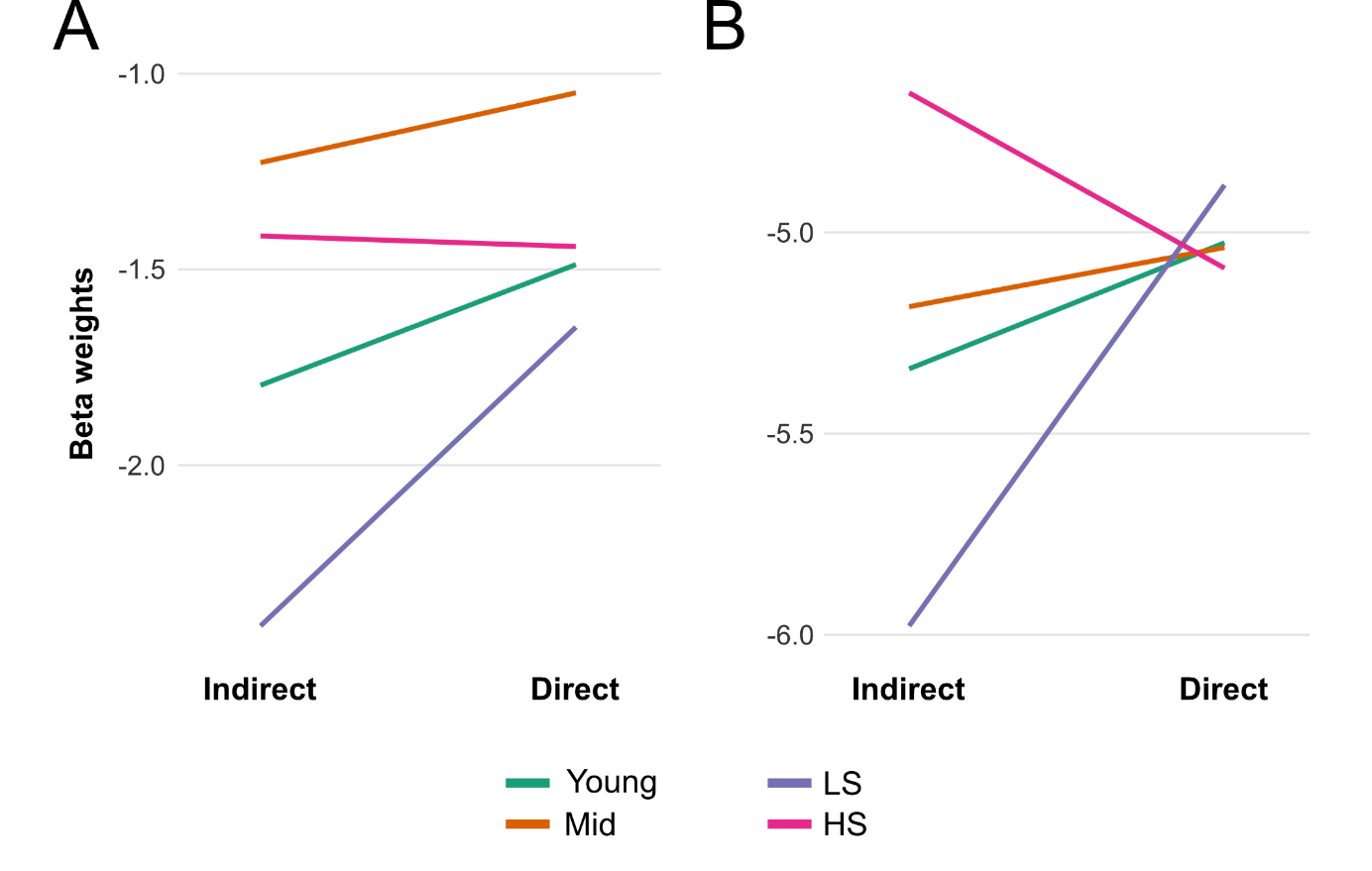

**Figure S9.** Mean beta weights per group and condition for (A) the cluster in the anterior/mid intraparietal sulcus and (B) the dorsal precuneal cluster (the ParPrec-clusters) from Bendtz et al. (2022) (see Figure S3 for anatomical location).

#### S3.1 Results from analyses using non-default procedures

We report the results from the Indirect > Direct and Direct > Indirect contrasts using non-default procedures (see §S2.5) in Tables S7 and S8, respectively. Significant clusters for the Indirect > Direct contrast in all adolescents and the Young and Mid groups are shown in Figure S10; results for the Direct > Indirect contrast in all adolescents and the Young group are shown in Figure S11. In general, we observed minor differences in the size of clusters previously identified using the default settings. However, two noteworthy differences emerged as additional clusters were found in both contrasts when pooling all adolescents. One of these clusters was also found in the Young group.

In the Indirect > Direct contrast, the non-default analysis yielded additional significant clusters in the bilateral orbitofrontal cortex/ventromedial PFC (vmPFC) for the Young group and all adolescents pooled. Posterior parts of these clusters showed activation under the default settings but were not significant (Young: *P* = .676, Adolescents: *P* = .900), markedly smaller (Young: k = 150, Adolescents: *P* = 107), and lay immediately at the anterior edge of the group masks used in the default second-level analysis. Notably, no significant cluster was observed in this region in Asaridou et al.’s (2019) study with adolescents using a similar contrast. However, the same area was activated in an “Affective > Informative” indirect speech act contrast (where the former were face-saving speech acts).

We also observed more extensive activation in the Indirect > Direct contrast within the anterior parts of the inferior temporal lobes.

Pooling all adolescents in the Direct > Indirect contrast yielded an additional significant cluster in the left anterior superior and middle frontal gyri. This region was also activated in the Young group in both the default and non-default procedures but appeared there as part of a larger cluster extending posteriorly into the triangular parts of the inferior frontal gyrus. An additional cluster was also revealed for the pooled adolescents in the right parahippocampal region.

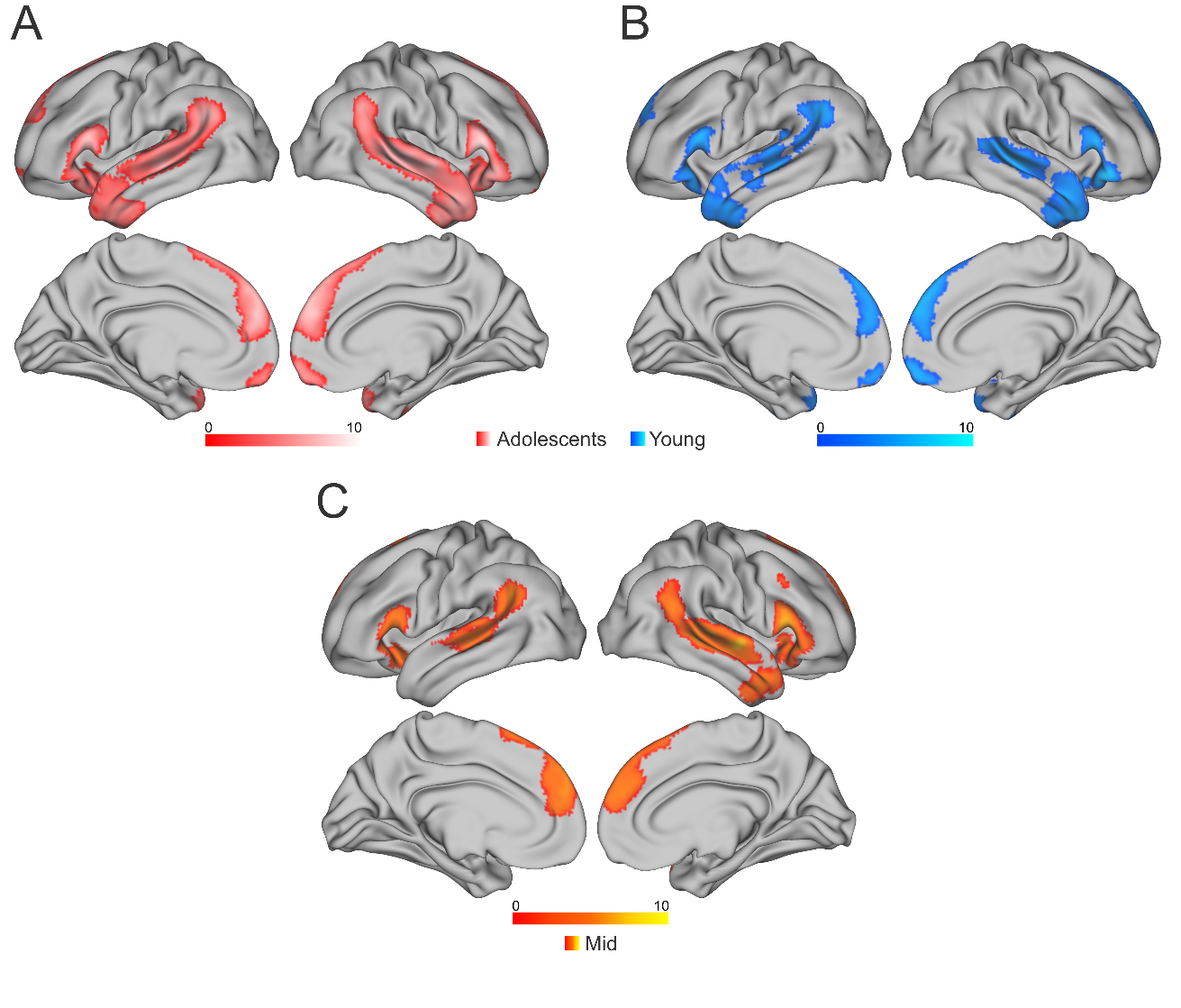

**Figure S10.** Significant clusters in the Indirect > Direct for all adolescents (A), the Young group (B), and the Mid group (C), using the non-default procedures described in section §S2.5. Significant clusters from the same contrast with default settings can be found in Figure S2A, 1A, and 1B, respectively. The non-default procedure revealed more extensive activation in ventral parts of the anterior temporal lobes. An additional significant cluster was revealed in the pooled adolescent group (A) and the Young group (B) in the bilateral anterior orbitofrontal/ventromedial prefrontal cortex (vmPFC).

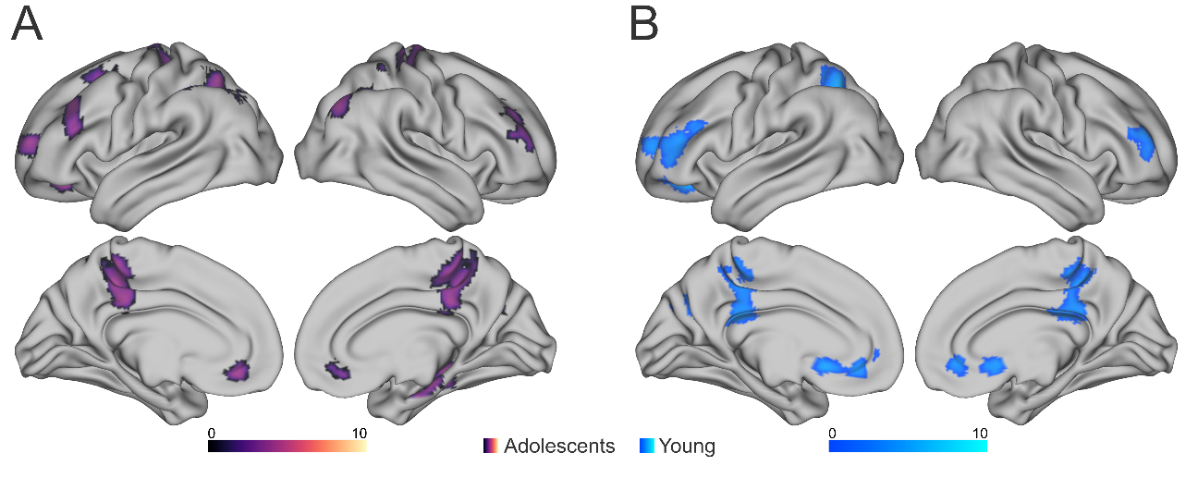

**Figure S11.** Significant clusters in the Direct > Indirect contrast for all adolescents (A) and the Young group (B), using the non-default procedures described in section §S2.5. Significant clusters from the same contrast with default settings can be found in Figure S2B and 2A, respectively. Pooling all adolescents together (A) using the non-default procedure revealed two additional clusters: the right parahippocampal gyrus and the left anterior middle and superior frontal gyri.

| Table S8. Significant clusters from the Indirect > Direct contrast for all adolescents and the Young and Mid group separately. These tests used the non-default procedures described in §S2.5. | | | | | | | | |
| --- | --- | --- | --- | --- | --- | --- | --- | --- |
| Anatomical region | **Local maxima (MNI)** | | | **Cluster** | | **Voxel** | | |
|  | x | y | z | k | *P*_FWE_ | *T*-value | *P*_FWE_ | |
| All adolescents |  |  |  |  |  | *T*(50) |  | |
| Right M/STG/ATL/IFG/AG | 58 | -34 | 2 | 7814 | < .001 | 8.76 | < .001 | |
| Bilateral mSFG/mPFC/anterior cingulate | -8 | 54 | 28 | 4016 | < .001 | 7.87 | < .001 | |
| Left M/STG/ATL/IFG/AG/SMG | -54 | 20 | 10 | 6832 | < .001 | 7.49 | < .001 | |
| Left cerebellum Crus I/II | -30 | -86 | -34 | 838 | < .001 | 5.90 | .016 | |
| Bilateral anterior orbitofrontal cortex/vmPFC | 6 | 58 | -16 | 496 | .009 | n.s. | | |
| Young |  |  |  |  |  | *T*(23) |  | |
| Right ATL/IFG | 50 | 32 | -8 | 2590 | < .001 | 7.26 | | . 018 |
| Bilateral anterior mSFG/mPFC | 8 | 58 | 26 | 2161 | < .001 | 7.01 | | .028 |
| Right M/STG | 56 | -34 | -2 | 1717 | < .001 | n.s. | | |
| Bilateral orbitofrontal cortex | 6 | 52 | -18 | 390 | .029 | n.s. | | |
| Left ATL/IFG | -60 | 18 | 18 | 2163 | < .001 | n.s. | | |
| Left posterior MTG/AG/SMG | -50 | -58 | 32 | 2137 | < .001 | n.s. | | |
| Mid |  |  |  |  |  | *T*(26) |  | |
| Right M/STG/ATL/IFG/AG | 62 | -10 | -4 | 4711 | < .001 | 8.75 | < .001 | |
| Left IFG/Insula | 24 | 20 | -12 | 954 | < .001 | 7.05 | .015 | |
| Left M/STG/AG/SMG | -52 | -26 | -2 | 1695 | < .001 | n.s. |  | |
| Bilateral anterior/posterior mSFG/SMA | -8 | 52 | 26 | 2770 | < .001 | n.s. |  | |
| Left Cerebelum Crus I/II | -20 | -76 | -28 | 794 | < .001 | n.s. |  | |

| Table S9. Significant clusters from the Direct > Indirect contrast for all adolescents and the Young and Mid group separately. These tests used the non-default procedures described in §S2.5. | | | | | | | |
| --- | --- | --- | --- | --- | --- | --- | --- |
| Anatomical region | **Local maxima (MNI)** | | | **Cluster** | | **Voxel** | |
|  | x | y | z | k | *P*_FWE_ | *T*-value | *P*_FWE_ |
| All adolescents |  |  |  |  |  | *T*(50) |  |
| Bilateral dPCC/precuneus/left anterior intraparietal sulcus | -26 | -24 | 30 | 3839 | < .001 | n.s. | |
| Left mid-to-posterior intraparietal sulcus/superior parietal lobule/middle occipital gyrus | -30 | -68 | 54 | 1051 | < .001 | n.s. | |
| Left posterior M/SFG | -28 | 22 | 60 | 471 | .012 | n.s. | |
| Mid-to-left posterior orbitofrontal cortex | -20 | 34 | -14 | 606 | .003 | n.s. | |
| Right parahippocampal gyrus/hippocampus/fusiform gyrus | 38 | -30 | -12 | 570 | .004 | n.s. | |
| Right AG/posterior intraparietal sulcus/middle occipital gyrus | 40 | -72 | 40 | 485 | .010 | n.s. | |
| Left anterior M/SFG | -26 | 62 | 12 | 390 | .032 | n.s. | |
| Left MFG/IFG (triangular part) | -36 | 32 | 36 | 664 | .001 | n.s. | |
| Right anterior MFG | 34 | 40 | 46 | 814 | < .001 | n.s. | |
| Right precentral and postcentral gyrus | 18 | -20 | 66 | 435 | .018 | n.s. | |
| Young |  |  |  |  |  | *T*(23) |  |
| Left M/SFG/IFG (triangular part)/posterior orbitofrontal cortex | -20 | 30 | -12 | 2219 | < .001 | n.s. |  |
| Left mid-to-posterior intraparietal sulcus/superior parietal lobule/precuneus | -32 | -62 | 54 | 1033 | < .001 | n.s. |  |
| Bilateral P/MCC/precuneus | 18 | -44 | 48 | 1496 | < .001 | n.s. |  |
| Right anterior MFG | 44 | 56 | 12 | 513 | .006 | n.s. |  |
| Mid |  |  |  |  |  |  |  |
| No significant clusters | | | | | | | |

#### S3.2 Analysis with only right-handed adolescents

The results from the one-sample tests (Adolescents, Young, and Mid) with only right-handed adolescents were consistent with the main analyses. Results are reported in Tables S9-10 and Figures S12-13. In the Indirect > Direct contrast with all adolescents pooled together, an additional cluster emerged in the precuneus/dPCC (Table S9, Figure S12A). This cluster does not overlap with the dPCC cluster from the HS > Adolescents contrast (Figure 4).

As reported in the article, we did not find any significant group differences between the two adolescent groups when running the whole-brain tests (Indirect > Direct, Young >/< Mid). When running the analysis with only right-handed participants, a significant cluster (*P_FWE_* < .029, k = 448) emerged in the left frontal lobe (Table S9 and Figure S14) in the direction Mid > Young. When overlaying the cluster with the AAL3 atlas (Rolls et al., 2020) in SPM, the cluster does however to a large part (~47%) not overlap with any specified region and instead spans white matter between, e.g., the left middle frontal gyrus and insula. Due to this lack of specificity as well as this cluster only emerging in our follow-up analysis to control for handedness, we refrain from interpreting this result any further.

##### S3.2.1 Additional analysis excluding adults with potential right-hemispheric language lateralization

As handedness information was unavailable for the adult participants, we conducted additional analyses based on hemispheric lateralization in the Context > ITI contrast (i.e., listening to speech > silence). Although this contrast was not designed as a language localizer task, it provides an approximate measure of lateralization in temporal auditory/language regions.

For each participant, we calculated a laterality index (LI; Seghier, 2008) based on the number of voxels in clusters in the left and right temporal lobes:

$$LI=\frac{Voxels_{Left}-Voxels_{Right}}{Voxels_{Left}+Voxels_{Right}}$$

Participants with an LI ≤ -0.2 were classified as showing potential right-hemispheric lateralization. This resulted in the exclusion of two participants from the LS group and four from the HS group. The four non-right-handed adolescent participants were also excluded, as in the additional analysis reported in S3.2.

The results were generally consistent with those in our main analysis and the whole-brain results are reported in Table SX. The *T*-tests in the dPCC followed a similar pattern to the main analysis (HS > LS: *T*(49) = 2.73, *P* = .004; LS > Adolescents: *T*(38) = 0.05, *P* = .52; LS > Adolescents (Bayesian): BF = 0.24). The non-significant ANOVAs in the amSFG and ATL remained non-significant (*F*(2, 95) = 0.52, *P* = .59, and *F*(2, 95) = 1.11, *P* = .33, respectively). The group effect in the pmSFG no longer reached statistical significance, *F*(2, 95) = 2.07, *P* = .13, compared with *F*(2, 105) = 3.62, *P* = .03 in the main analysis. Despite the reduction in sample size by ten participants, the results were generally consistent with the main analysis. We therefore do not consider the outcome of this additional analysis to alter our overall conclusions.

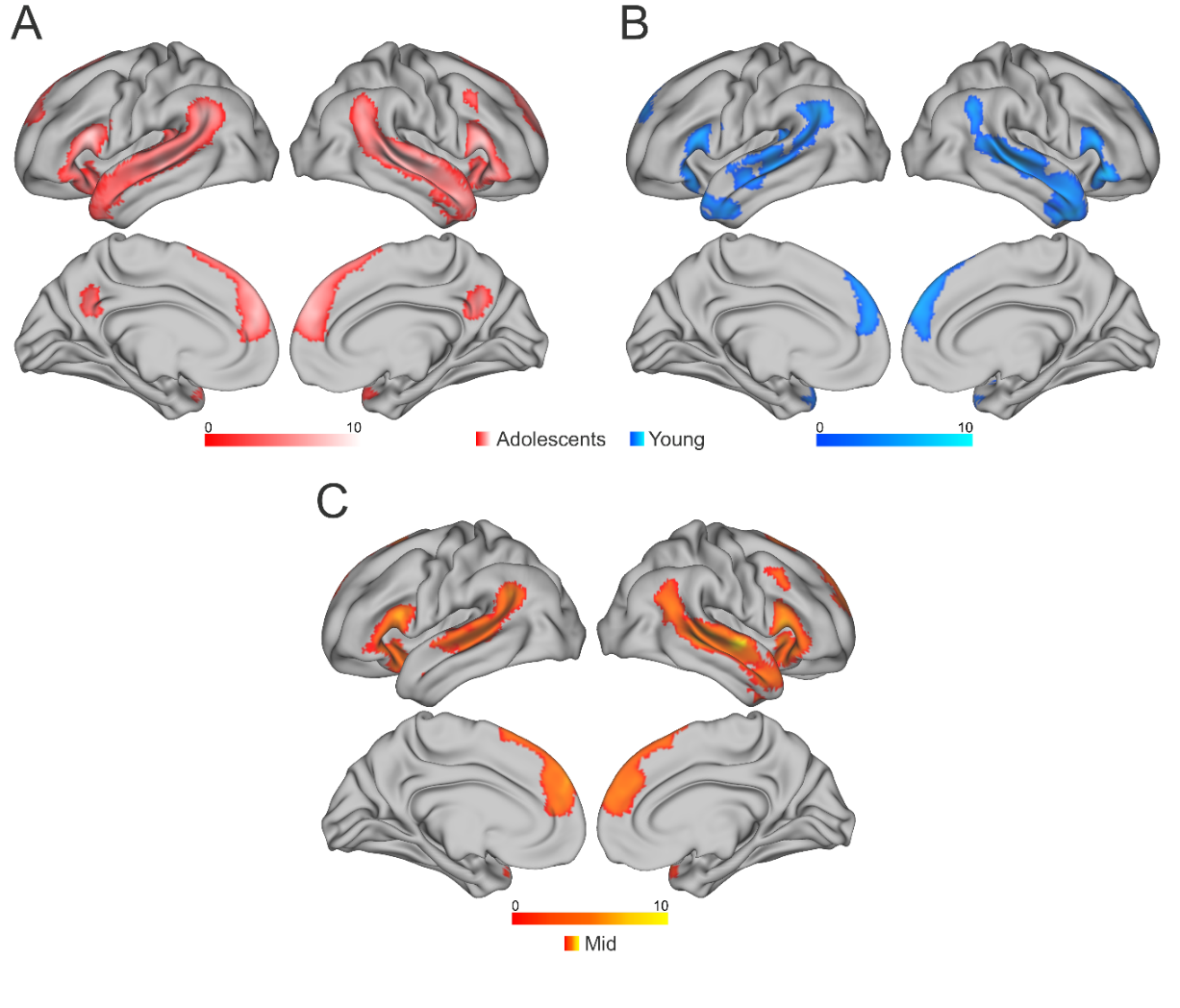

Figure S12. Significant clusters in the Indirect > Direct for all adolescents (A), the Young group (B), and the Mid group (C), with only right-handed participants. The results are consistent with the analysis with all participants and do not change the interpretations of our main result. A significant cluster appeared in the precuneus for all adolescents (A) but did not overlap with the dPCC cluster (Figure 4).

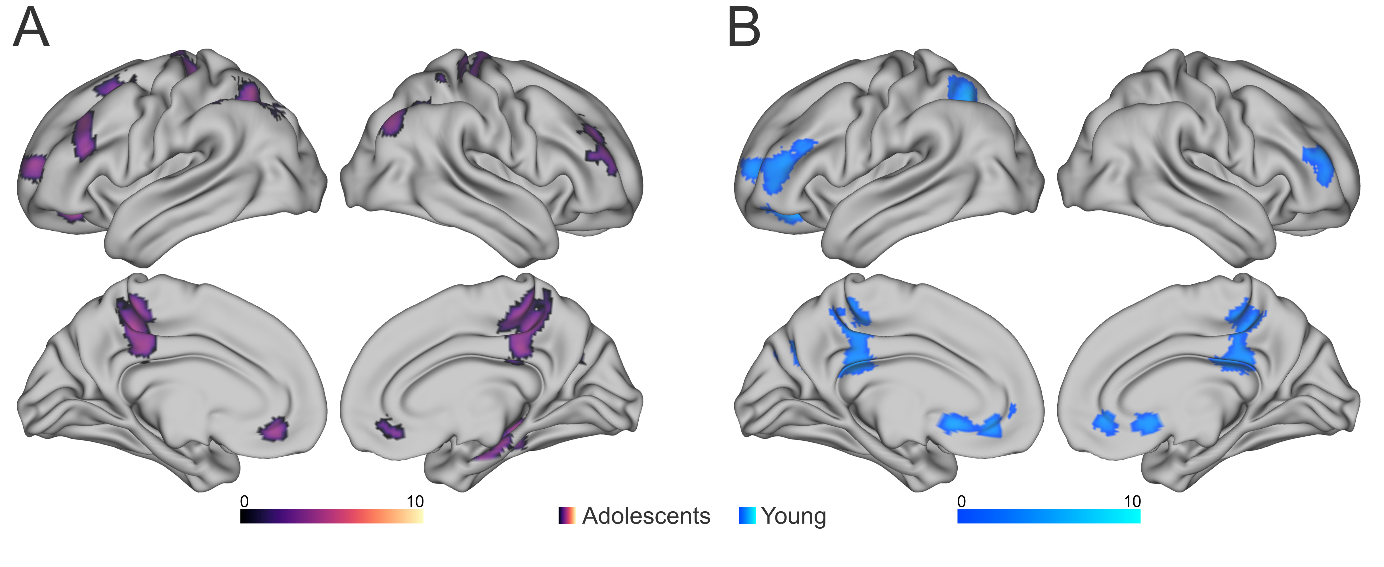

Figure S13. Significant clusters in the Direct > Indirect contrast for all adolescents (A) and the Young group (B), with only right-handed participants. The results are consistent with the analysis with all participants and do not change the interpretations of our main result

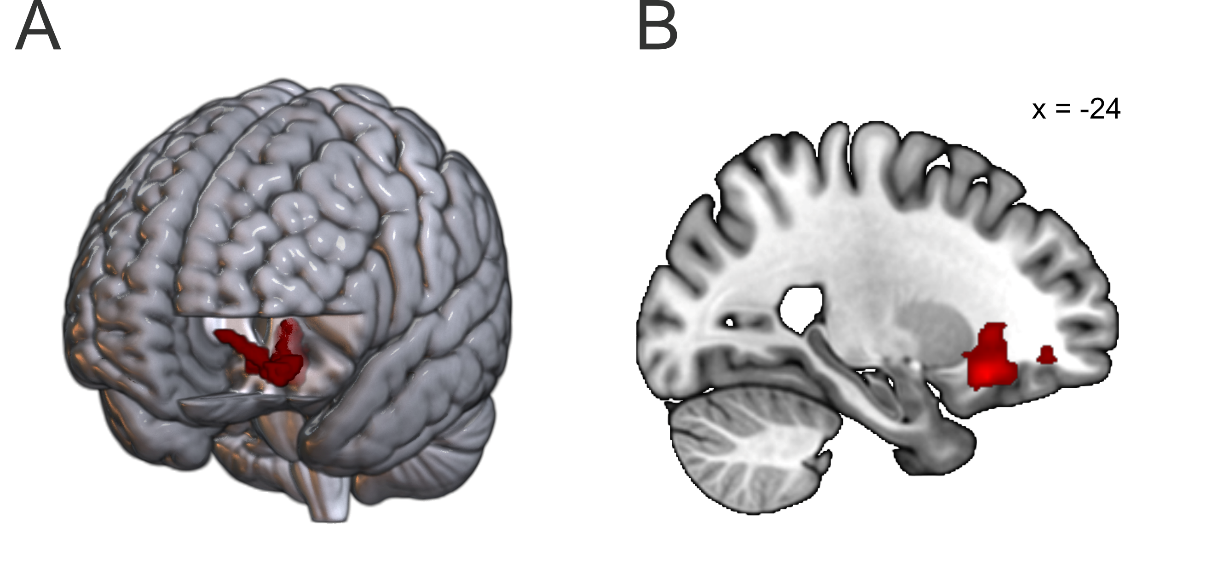

**Figure S14.** Significant cluster in the Indirect > Direct, Mid > Young contrast with only right-handed participants. The cluster was located in the left frontal lobe. ~47% of the cluster did not overlap with any specified region in AAL3 (Rolls et al., 2020). (B) shows cluster in a sagittal slice at MNI coordinate x = -24.

| Table S10. Significant clusters from the Indirect > Direct contrast for all adolescents and the Young and Mid group separately with only right-handed participants. Significant clusters are also reported from tests for group effects between the two groups. | | | | | | | |
| --- | --- | --- | --- | --- | --- | --- | --- |
| Anatomical region | **Local maxima (MNI)** | | | **Cluster** | | **Voxel** | |
|  | x | y | z | k | *P*_FWE_ | *T*-value | *P*_FWE_ |
| All adolescents |  |  |  |  |  | *T*(46) |  |
| Right M/STG/ATL/IFG/AG | 58 | -34 | 2 | 6722 | < .001 | 8.96 | < .001 |
| Left M/STG/IFG/AG | -54 | 20 | 14 | 6022 | < .001 | 7.78 | < .001 |
| Bilateral mSFG/mPFC/SMA/anterior cingulate | -8 | 54 | 28 | 4019 | < .001 | 7.63 | < .001 |
| Left cerebellum Crus I/II | -20 | -80 | -34 | 588 | .007 | 6.24 | .005 |
| Left precuneus/dPCC | 8 | -56 | 34 | 424 | .036 | 5.83 | .017 |
| Young > Mid | | | | | | | |
| No significant clusters | | | | | | | |
| Mid > Young | | | | | | *T*(45) |  |
| Left MFG/Insula | -24 | 22 | -14 | 438 | .029 | 5.49 | .047 |
| Young | | | | | | *T*(21) |  |
| Right M/STG/ATL/IFG/AG | 50 | -54 | 32 | 3874 | < .001 | 7.02 | .032 |
| Bilateral anterior mSFG/mPFC | 8 | 58 | 28 | 1848 | < .001 | n.s. |  |
| Left M/STG/AG/SMG | -60 | -50 | 14 | 2352 | < .001 | n.s. |  |
| Left IFG/ATL/I/MTG | -60 | 18 | 18 | 1223 | < .001 | n.s. |  |
| Mid | | | | | | *T*(24) |  |
| Right M/STG/IFG/AG/ATL/Insula | 62 | -6 | -4 | 4972 | < .001 | 11.25 | < .001 |
| Left Cerebellum Crus I/II | -22 | -78 | -36 | 994 | < .001 | 6.92 | .021 |
| Left IFG/Insula | -24 | 20 | -12 | 1391 | < .001 | 6.58 | .041 |
| Bilateral anterior/posterior mSFG/mpFC/SMA | -2 | 58 | 32 | 3228 | < -001 | n.s. | |
| Left M/STG/AG/SMG | -52 | -26 | -2 | 1984 | < .001 | n.s. | |

| Table S11. Significant clusters from the Direct > Indirect contrast for all adolescents and the Young and Mid group separately with non-right-handed participants removed. | | | | | | | |
| --- | --- | --- | --- | --- | --- | --- | --- |
| Anatomical region | **Local maxima (MNI)** | | | **Cluster** | | **Voxel** | |
|  | x | y | z | k | *P*_FWE_ | *T*-value | *P*_FWE_ |
| All adolescents |  |  |  |  |  | *T*(46) |  |
| Bilateral dPCC/precuneus | 30 | -10 | 36 | 2887 | < .001 | n.s. | |
| Left posterior M/SFG | -28 | 14 | 56 | 393 | .050 | n.s. | |
| Left mid-to-posterior intraparietal sulcus | -30 | -68 | 54 | 616 | .005 | n.s. | |
| Left M/SFG/IFG (triangular part) | -28 | 40 | 48 | 418 | .038 | n.s. | |
| Right anterior MFG | 38 | 56 | 12 | 468 | .022 | n.s. | |
| Young |  |  |  |  |  | *T*(23) |  |
| Left M/SFG/IFG/orbitofrontal cortex | -20 | 30 | -12 | 1610 | < .001 | n.s. | |
| Right MFG | 44 | 56 | 12 | 472 | .019 | n.s. | |
| Left mid-to-posterior intraparietal sulcus/precuneus | -32 | -62 | 54 | 879 | < .001 | n.s. | |
| Bilaterial dPCC/Precuneus | 18 | -44 | 48 | 1159 | < .001 | n.s. | |
| Mid |  |  |  |  |  |  |  |
| No significant clusters | | | | | | | |

| Table S12. Significant clusters from tests involving adult participants. Non-right-handed adolescents and adults with right lateralized language processing were excluded. | | | | | | | |
| --- | --- | --- | --- | --- | --- | --- | --- |
| Anatomical region | **Local maxima (MNI)** | | | **Cluster** | | **Voxel** | |
|  | x | y | z | k | *P*_FWE_ | *T*-value | *P*_FWE_ |
| Adults, Direct > Indirect |  |  |  |  |  | *T*(50) |  |
| Bilateral precuneus/intraparietal sulcus/superior parietal lobe | -28 | -66 | 40 | 8839 | < .001 | 6.34 | .004 |
| Bilateral middle and superior frontal gyrus | -24 | 42 | -10 | 9551 | < .001 | 5.85 | .018 |
| Left posterior inferior temporal gyrus/Cerebellum Crus I | -52 | -38 | -18 | 2139 | < .001 | n.s. | |
| Right Cerebellum Crus I/II/VIIb | 36 | -74 | -52 | 495 | .011 | n.s. | |
| LS, Direct > Indirect |  |  |  |  |  | *T*(25) |  |
| Bilateral dPCC/precuneus/intraparietal sulcus | -28 | -62 | 42 | 9090 | < .001 | 7.82 | .004 |
| Left posterior inferior temporal gyrus/Cerebellum Crus I/II/VI/VIII | -40 | -42 | -30 | 2450 | < .001 | 7.02 | .019 |
| Left middle and superior frontal gyrus | -24 | 40 | -10 | 5403 | < .001 | 6.84 | .026 |
| Right middle and superior frontal gyrus | 28 | 50 | 4 | 4642 | < .001 | n.s. | |
| Right Cerebellum Crus I/II/VIII | 30 | -36 | -38 | 1481 | < .001 | n.s. | |
| Right ventral PCC/cuneus | 16 | -52 | 20 | 404 | .019 | n.s. | |
| HS, Direct > Indirect |  |  |  |  |  |  |  |
| No significant clusters | | | | | | | |
| Old > Adolescents, Indirect > Direct |  |  |  |  |  |  |  |
| No significant clusters | | | | | | | |
| Adolescents > Old, Indirect > Direct |  |  |  |  |  |  |  |
| No significant clusters | | | | | | | |
| HS > Adolescents, Indirect > Direct |  |  |  |  |  | *T*(70) |  |
| Dorsal posterior cingulate cortex | 0 | -26 | 24 | 988 | < .001 | 5.23 | .037 |
| Adolescents > HS, Indirect > Direct |  |  |  |  |  |  |  |
| No significant clusters | | | | | | | |

| Table S13. Overlaps between ROIs (pmSFG, dPCC, amSFG, and ATL) and the ParPrec clusters from Bendtz et al. (2022) (treated separately) and atlas from (Du et al., 2024) (53 % agreement). Percentage of overlapping voxels is based on the number of voxels inside atlas, rather than the original cluster. For the dPCC, we used the clusters that came out of the analysis with default settings (see Table S2-3). | | | | | | | |
| --- | --- | --- | --- | --- | --- | --- | --- |
|  |  | **pmSFG** | **dPCC** | **amSFG** | **ATL** | **Par** | **Prec** |
|  | Cluster size | 956 | 1047 | 901 | 433 | 368 | 399 |
|  | Inside atlas | 49% (560) | 67% (700) | 63% (568) | 82% (354) | 74% (271) | 40% (161) |
| Networks | AUD |  |  |  | 8.5% |  |  |
|  | CG-OP | 4.6% | 1.6% |  |  |  |  |
|  | dATN-A |  |  |  |  | 34.7% | 50.3% |
|  | dATN-B |  |  |  |  | 0.7% |  |
|  | DN-A |  | 26.0% |  |  |  |  |
|  | DN-B | 13.4% | 10.9% | 94.5% | 54.5% |  |  |
|  | FPN-A | 0.2% | 2.6% |  |  | 47.6% | 16.1% |
|  | FPN-B | 0.5% | 10.1% | 1.4% |  | 3.7% |  |
|  | LANG | 80.7% |  | 4.0% | 45.5% |  |  |
|  | PM-PPR |  |  |  |  | 12.5% |  |
|  | SAL/PMN | 0.5% | 48.0% |  |  |  | 33.5% |
|  | SMOT-A |  | 0.9% |  |  | 0.7% |  |
| *Note: Abbreviations of networks (Du et al., 2024): AUD: Auditory; CG-OP: Cingulo-Opercular; dATN-A: dorsal Attention Network A; dATN-B: dorsal Attention Network B; DN-A: Default Network A; DN-B: Default network B; FPN-A: Fronto-Parietal Network A; FPN-B: Fronto-Parietal Network B; LANG: Language; PM-PPR: Premotor-Posterior Parietal Rostral; SAL/PMN: Salience/Parietal Memory Network; SMOT-A: Somatomotor A.* | | | | | | | |

### S4. Supplementary discussion

#### S4.1. Activity in the anterior and posterior medial superior frontal gyrus

While we had hypothesized that the anterior parts of the medial superior frontal gyrus (amSFG) might show decreased activity for ISA, at least from adolescence into adulthood, we did not observe any significant age-related differences, although all groups activated this region during ISA processing. This suggests that the increase in posterior (pmSFG) activity is not necessarily a replacement of anterior processes, but an additional resource recruited when interpreting indirect speech acts. This is consistent with our view that the anterior portion of this structure and the anterior medial prefrontal cortex may govern general purpose/abstract ToM processes, relative to the more context-dependent posterior aspect. The results indicate that these processing types happen in parallel, at least in more mature individuals, during the understanding of indirect speech acts.

Contextualizing the activity observed in the pmSFG within further neuropragmatic work, the cluster shows some overlap with one of six clusters from a recent meta-analysis of non-literal language comprehension (including ISA, Hauptman et al., 2023), most of which covered language and ToM areas. Our proposal outlined in the main manuscript of a potential function for this cluster in the context of pragmatic comprehension is thus consistent with other present literature generalizing over several pragmatic (non-literal) phenomena.

#### S4.2 Supplemental finding in the ventral medial prefrontal/orbitofrontal cortex

Out of the supplementary analysis using non-default masking settings (§S2.5 and §S3.1), we wish to report one additional cluster. The cluster appeared for the Young group and all adolescents pooled together in the Indirect > Direct contrast and was located in the anterior parts of the ventromedial prefrontal/orbitofrontal cortex (referred to here as vmPFC, see Figure S11). This cluster did not appear in the main analysis due to poor voxel coverage within this region. While this supplementary finding does not alter our main interpretations, we note that a similar cluster was present among adolescents in a face-saving > non-face-saving indirect speech act contrast in Asaridou et al.’s (2019) (unpublished) study. The vmPFC is considered part of the ToM network, though more associated with affective than cognitive ToM (the latter is rather supported by more dorsally located prefrontal locations; Healey & Grossman, 2018; Sebastian et al., 2012; Shamay-Tsoory et al., 2006). We therefore suggest that the activity in the vmPFC may reflect the affective/face-saving dimension of indirect criticisms and rejections—i.e. processing the emotional aspects of the conversation.
